# Dynamic Network Analysis of Electrophysiological Task Data

**DOI:** 10.1101/2024.01.12.567026

**Authors:** Chetan Gohil, Oliver Kohl, Rukuang Huang, Mats W.J. van Es, Oiwi Parker Jones, Laurence T. Hunt, Andrew J. Quinn, Mark W. Woolrich

## Abstract

An important approach for studying the human brain is to use functional neuroimaging combined with a task. In electrophysiological data this often involves a time-frequency analysis, in which recorded brain activity is time-frequency transformed and epoched around task events of interest, followed by trial-averaging of the power. Whilst this simple approach can reveal fast oscillatory dynamics, the brain regions are analysed one at a time. This causes difficulties for interpretation and a debilitating number of multiple comparisons. In addition, it is now recognised that the brain responds to tasks through the coordinated activity of networks of brain areas. As such, techniques that take a whole-brain network perspective are needed. Here, we show how the oscillatory task responses from conventional time-frequency approaches, can be represented more parsimoniously at the network level using two state-of-the-art methods: the HMM (Hidden Markov Model) and DyNeMo (Dynamic Network Modes). Both methods reveal frequency-resolved networks of oscillatory activity with millisecond resolution. Comparing DyNeMo, HMM and traditional oscillatory response analysis, we show DyNeMo can identify task activations/deactivations that the other approaches fail to detect. DyNeMo offers a powerful new method for analysing task data from the perspective of dynamic brain networks.

**Highlights:**

- We show how oscillatory task response analysis can be carried out at the network level using two state-of-the-art methods: the Hidden Markov Model (HMM) and Dynamics Network Modes (DyNeMo).
- The HMM and DyNeMo can identify oscillatory task responses that conventional time-frequency methods fail to detect.
- DyNeMo provides a more interpretable and precise network decomposition of the data compared to the HMM.
- The dataset and (Python) scripts used to perform dynamic network analysis on task data are made publicly available.

## 1 Introduction

Non-invasive neuroimaging techniques, such as functional magnetic resonance imaging and magneto/electroencephalography (M/EEG), provide a means for viewing the human brain during function. These imaging techniques are often used in a *task paradigm*. Here, a participant is asked to perform a particular task while their brain activity is recorded^1^. The task is designed to isolate a particular cognitive process, thereby studying the brain activity in response to the task can reveal insights into how the brain performs the necessary cognition for the task.

A well known property of the brain is the emergence of oscillatory activity from neuronal populations, which can occur both at rest and in response to a task [1]. M/EEG is a particularly useful modality for studying these oscillations because of its high temporal resolution and the fact that it is a direct measure of neuronal activity.

A very popular method for analysing task M/EEG data is to compute oscillatory responses using time-frequency (TF) decompositions [3]. This approach looks at the trial-averaged, oscillatory response that occurs in a task at a single spatial location, e.g. a brain region. However, such an approach neglects the interaction of a particular region with the rest of the brain.

More recently, the widespread adoption of large-scale neural recordings and whole-brain imaging techniques has prompted neuroscientists to develop new frameworks for understanding brain activity at the level of inter-regional interactions, rather than focusing solely on region-specific responses [4, 5]. A potentially more powerful approach for studying M/EEG data would be to take a large-scale network perspective. After all, it is widely believed that the human brain performs cognition via large-scale functional networks [6]. Therefore, identifying the functional networks that activate in response to a task may yield a better understanding of the cognitive processes occurring in the brain. This also alleviates the burden of doing separate analyses on each region, in terms of both the difficulties of interpretation and the multiple comparison correction.

A popular hypothesis for the mechanism of communication in the brain is via the phase locking of oscillatory activity in different neuronal populations (‘communication via coherence’ [7]). Given this hypothesis, we expect the brain to exhibit large-scale functional networks of coherent activity. Indeed, this has been shown to be the case, fast transient networks of coherent activity have been identified in task [8] and at rest [9]. Both [8] and [9] used an HMM (Hidden Markov Model) [10] to identify transient networks in M/EEG data. This is an *unsupervised* method, meaning the model learns the timing of network activations directly from the data without any input from the user.

An important assumption made by the HMM is that only one network is active at a given time point. More recently, a more flexible unsupervised dynamic network model called DyNeMo (Dynamic Networks Modes) has been introduced [11]. Crucially, this model allows co-activations of multiple networks to occur. Multiple cognitive processes in the brain are likely to occur simultaneously. Each cognitive process may be underpinned by its own large-scale functional network [6]. Therefore, we hypothesise that DyNeMo could provide a better model for understanding cognitive processes in the brain, and could be particularly useful in the context of task data analysis.

Here, we show how we can use the network descriptions of the HMM and DyNeMo to analyse task M/EEG data. This can be thought of as doing a task response analysis at the network level rather than computing event-related responses for the activity in every brain area separately. We compare conventional TF methods for computing oscillatory responses when analysing task data, to the dynamic network-level approaches of the HMM and DyNeMo. Using simulations with overlapping network activations, we that show DyNeMo can identify co-activations/deactivations that the HMM cannot. Using a publicly available real MEG task dataset, we show qualitatively that the network-based perspective provided by the HMM and DyNeMo can identify task responses that conventional oscillatory response methods fail to detect. Finally, we show that DyNeMo can provide a more precise dynamic network description compared to the HMM, due to its ability to infer brain networks with temporally overlapping dynamics.

## 2 Methods

### 2.1 Datasets

#### 2.1.1 Simulation Data

To evaluate the performance of the HMM and DyNeMo in identifying oscillatory networks, we trained both models on simulated data containing bursting oscillatory activity. Bursts were simulated using a two-state Markov chain with the transition probability matrix

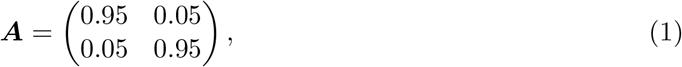

where one state corresponds to an ‘on’ state and the other an ‘off’ state. The burst time course for each network was independently sampled from this transition probability matrix. The observed data was simulated at each region of interest (parcel) in source space as

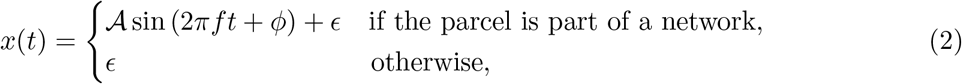

where 𝒜 is the amplitude, *f* is the oscillatory frequency of the network, *t* is time, *ϕ* ∼ Uniform(0, 2*π*) is a random phase, and *ϵ* ∼ 𝒩 (0, *σ*^2^) is Gaussian noise with variance *σ*^2^ = 0.2. A random phase lag was simulated between regions and symmetric orthogonalisation [12] was applied to remove zero-lag correlations. At a sampling frequency of 250 Hz, with the transition probability matrix in Eq. (1), this leads to an average lifetime of approximately 40 ms. The amplitude of the bursts alternated between a value of 𝒜 = 1 for low-amplitude events and 𝒜 = 2 for high-amplitude events.

Two networks were simulated: a visual network with activity at *f* = 10 Hz in six parcels in the occipital lobe and a motor network with activity at *f* = 20 Hz in four parcels in the motor cortex. There were 38 parcels in total and we simulated 25600 time points. We applied time-delay embedding with *±*2 lags^2^ and principal component analysis (PCA) to reduce the dimensionality down to 120 channels (approximately 85% explained variance), see Section 2.3 for further details. Finally, we standardised (z-transform) the data before training a model.^3^

#### 2.1.2 Real Task MEG Data

In this report, we will study a simple visual perception task. 19 participants (11 male, 8 female, 23-37 years old) were presented with an image of famous or unfamiliar face or a scrambled image. The participants were asked to judge the symmetry of the image via a button press. This button press was included to ensure the participant focused on the images presented. Figure 1 shows a schematic of a single trial. Each participant completed 6 sessions^4^ that contained approximately 200 trials evenly split between famous, unfamiliar and scrambled images. See [19] for further details regarding the experimental paradigm and dataset.

**Figure 1.**
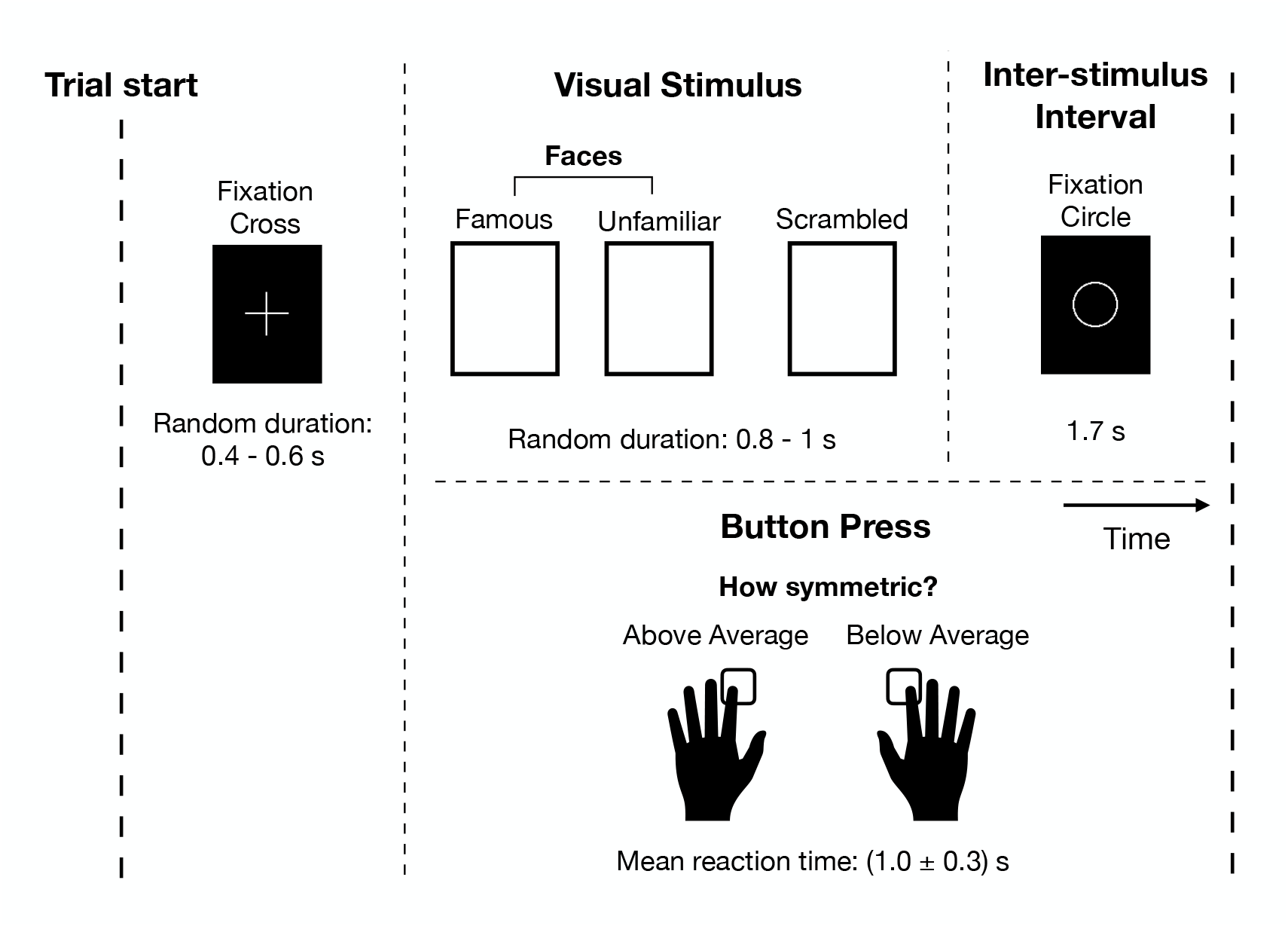
Single-trial schematic of the task. Participants were presented with an image of a face (famous or unfamiliar) or a scrambled image and pressed a button in response to indicate their judgement on the symmetry of the image. See [19] for further details.

The public dataset provides the MaxFiltered data. The following steps were applied to the data using the OHBA Software Library (OSL) [20]:

##### Preprocessing

Starting from the MaxFiltered data, we apply some basic preprocessing. This includes filtering the data, downsampling and automated artefact removal. The exact preprocessing steps we applied are summarised in supplementary information (SI) Figure S3A.

##### Source reconstruction

Starting from the preprocessed sensor-level data, we perform source reconstruction using a Linearly Constrained Minimum Variance (LCMV) beamformer [21]. This involves projecting the preprocessed sensor data onto a 8 mm dipole grid inside the inner skull. We use the brain/skull surfaces extracted from each subject’s structural MRI for the head model. The noise covariance matrix used to calculate the beamformer was estimated using the sensor-level data for a subject and regularised to a rank of 60 using PCA. Note, the MaxFiltering step reduces the rank of the sensor-level data to approximately 64 and there is an additional reduction in the rank between 1-4 due to artefact removal with independent component analysis (ICA). This resulted in the choice of 60 for the rank of the sensor-level data covariance. We use a unit-noise-gain invariant LCMV beamformer [22], which normalises the beamformer weights to ensure depth does not impact the amplitude of source signals.

##### Parcellation

Using the dipole time courses from source reconstruction, parcel time courses were calculated using a 38 region of interest (non-binary, probabilistic) parcellation. First, the dipole time courses are weighted (multiplied) by the probability of belonging to a parcel. Then, PCA is performed on the weighted dipole time courses for each parcel and we take the first principal component as the parcel time course. Following this, we apply symmetric orthogonalisation to the parcel time courses [12] to minimise spatial leakage and dipole sign flipping to align the sign of each parcel time course across subjects. The exact steps we applied are summarised in SI Figure S3B.

### 2.2 Conventional Single-Channel Methods

#### TF response

A common approach for studying task MEG data is to compute oscillatory responses using TF transformations [3]. While the ordering of steps can somewhat vary, this involves calculating the TF response as shown in Figure 2: first, the continuous data is epoched around the occurrence of an event and a TF transformation (typically a wavelet [3]) is calculated using the epoched data and the absolute value is taken. Finally, we trial average to give the full TF response. The full TF oscillatory response contains both the fixed latency (task-phase-locked) response, known as the *evoked response*, and the jittered latency (non-task-phase-locked) response, known as the *induced response* [14]. We can estimate the evoked TF response by TF transforming the trial-averaged time series rather than each trial individually. Subtracting the evoked TF response from the full TF response, we can estimate the induced TF response. This calculation can be done at the sensor (outside of the brain) or source (within the brain) level. The TF response is calculated for each channel separately. Note that in this paper, channels can correspond to sensors or to parcels in brain space. The result of such an analysis is a 2D array (time by frequency) for the evoked/induced TF response for each subject and channel.

**Figure 2.**
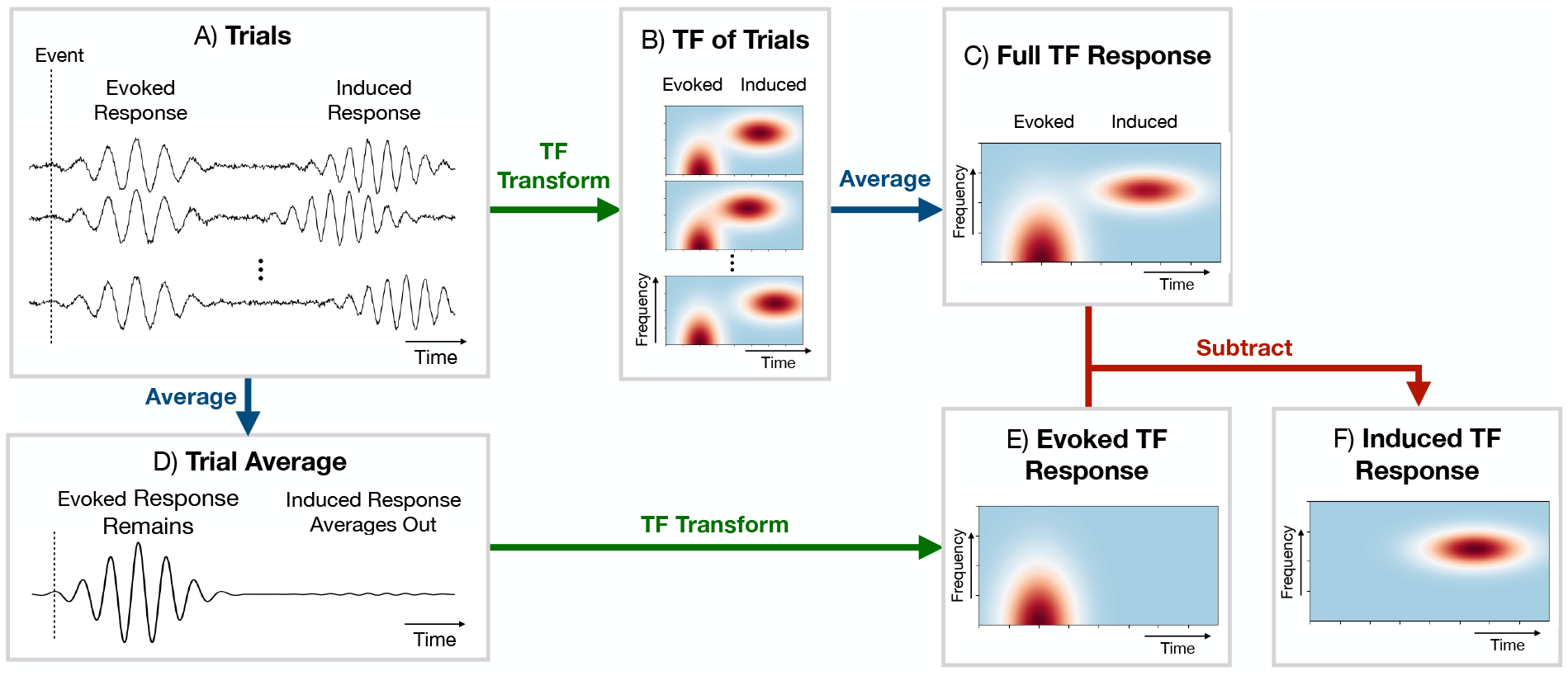
Calculation of a time-frequency (TF) response in conventional single-channel analyses. A) Individual trials (calculated by epoching the data around a particular event). The individual trials can contain both an evoked (phase-locked to the task, i.e. fixed latency) and induced (not phase-locked to the task, i.e. jittered latency) response. B) TF transform and calculation of oscillatory power at each time and frequency (e.g. a wavelet transform) of each trial. C) Full TF oscillatory response calculated by averaging the trial-specific TF responses; note that this contains both the evoked and induced response. D) Average response over the trials of the data prior to the TF transform. E) Evoked TF response calculated by TF transforming the trial average from (D). F) Induced TF response calculated by subtracting the evoked TF response from the full TF response.

In this paper, we will epoch the data using a window from *t* = −100 ms to 1 s, where *t* = 0 s is when an event occurs and *t* is time. We will use a Morlet wavelet [3] with a length of four cycles to calculate the TF transformation for the frequency range 6-30 Hz using a frequency bin width of 0.5 Hz. To reduce memory requirements, we will slide the Morlet wavelet with a spacing of three samples, i.e. use a decimation factor of three, which lowers the temporal resolution of TF transformation by a factor of three.

Often it is necessary to *baseline correct* the TF response to isolate the change in activity due to the event [3]. In this paper, we will subtract the mean TF response from *t* = −100 ms to 0 s for each frequency separately. This is the last step after calculating the evoked/induced TF response.

#### Statistical significance testing

Once we have calculated a TF response for each subject and channel, we need to test for statistical significance to check whether the response we have observed can be simply due to chance. We test if the average TF response across subjects (*group average*) is significant for each time point and channel. The null hypothesis is that the event does not evoke/induce a TF response, i.e. a TF response of zero. We calculate a *p*-value for the TF response we have observed given this null hypothesis and if the response is sufficiently unlikely (has a *p*-value *<* 0.05), we conclude it is significant. In this report, we use non-parametric permutation testing with a GLM (General Linear Model) [15] to do this. We use the maximum COPE (Contrast of Parameter Estimate) across time and channels to control for multiple comparisons (familywise error rate). See SI Section 1.1 for further details.

### 2.3 Dynamic Network Models and Post-Hoc Analysis

Here, we introduce the two state-of-the-art methods we will use for identifying dynamic networks in the task MEG data: the HMM [16] and DyNeMo [11]. Both methods infer networks without any knowledge of the task. We used the OHBA Software Library Dynamics toolbox (osl-dynamics) [18] implementation of these models in this work.

#### HMM

Here, we use an HMM with a multivariate normal observation model. This model assumes the data is generated by a finite set of *states*, each with its own characteristic covariance matrix (which represents a network). We assume the mean vector is zero. Only one state may be active at a particular time point. Based on this, the model learns a time course for the probability of a state being active at each time point in the input data. Both the state covariance matrices (i.e. functional networks) and probabilities are learnt directly from the data, without any input from the user. We use the “TDE-HMM” variant of this model, which was introduced in [9]. See [18] for further details describing this generative model.

#### DyNeMo [11]

Here, we also use a multivariate normal observation model. However, DyNeMo assumes the data is generated by a set of *modes* [11], each with its own characteristic covariance matrix (assuming zero mean). Crucially, these modes can overlap. The model learns the coefficient for how much each mode contributes to the total (instantaneous) covariance of the data at a given time point^5^. In Section 3.1, we will show this model outperforms the HMM using simulated data when the ground truth includes co-activating network bursts. Both the mode covariance matrices (i.e. functional networks) and mode time courses are learnt directly from the data, without any input from the user. See [11, 18] for a detailed description of DyNeMo and a comparison with the HMM. SI Figure S1 compares the generative model for DyNeMo to the HMM.

#### Time-delay embedding

Before training each model, we prepare the data by performing time-delay embedding [18]. This is a process of augmenting the data with additional channels that contain time-lagged versions of the original channels. By doing this, we encode the spectral properties of the original data into the covariance matrix of the time-delay embedded data. See SI Figure S2 for description of how this works. This step allows us to model frequency-specific phase coupling (coherence) between pairs of parcels (as encoded by the cross-correlation function into the covariance matrix). Note, phase coupling between parcels can only occur at a particular frequency here. In DyNeMo, cross-frequency coupling would manifest as a co-activation of modes with activity at different frequencies.

In this paper, we will embed 14 additional channels for each original channel corresponding to lags of -7 to 7 samples. This corresponds to an embedding window of *±*30 ms with 250 Hz data. We will also reproduce our analysis using a different number of lags to ensure our results are robust to this choice (see SI Figure S8). After time-delay embedding, we’re often left with a very high-dimensional time series. We will use PCA to reduce the number of channels to 80 to reduce the computational load in training a model. 80 PCA components was chosen because it explained a high amount of variance (approximate 70%) in the time-delay embedded data. The final step before training a model is to standardise (z-transform) the data.

The data preparation and dynamic network modelling is summarised in SI Figure S3C. The use of *±*7 lags in time-delay embedding is a good choice for 250 Hz data for researchers that are interested in 1-45 Hz activity in the original data. We recommend using enough PCA components to explain approximately 70% of variance.

#### Post-hoc spectra

After training the HMM/DyNeMo, we calculate state/mode spectra using the inferred state/mode time courses and the source (parcel) data. For the HMM, we use the multitaper approach described in [16], and for DyNeMo, we use the GLM-spectrum approach described in the SI of [11]. Both of these methods provide high-resolution power/cross spectral densities (P/CSDs) for each parcel and pair of parcels for each state/mode and subject. **Power maps**. We calculate power maps from the group-average state/mode PSDs. Power maps are calculated by taking the average value across a frequency band in the PSD for each parcel. For the HMM, we use the non-negative matrix factorisation (NNMF) approach introduced in [9] to select the frequency range. We take the first NNMF component from fitting two components to the stacked coherence spectra from each subject and state. This typically covers the frequency range 1-22 Hz. For DyNeMo, we average the mode PSDs across the entire frequency range (1-45 Hz). When visualising the power maps, we will show them relative to mean across states/modes without any thresholding.

#### Coherence networks

We use the coherence averaged over a frequency range as our measure of functional connectivity. To calculate coherence networks we first calculate the coherency spectrum from the group-average P/CSDs using

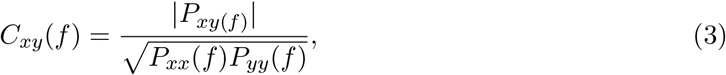

where *P*_*xy*_(*f*) is the CSD for parcels *x* and *y*, and *P*_*xx*_(*f*) (*P*_*yy*_(*f*)) is the PSD for parcel *x* (*y*). We then take the average value across a frequency range to give a single value for the coherence for the edge connecting parcel *x* and *y*. For the HMM, we use the first NNMF component (same as the power maps, approximately 1-22 Hz) to specify the frequency range. For DyNeMo, we use the full frequency range (1-45 Hz). For visualisation, we show the HMM coherence network relative to the mean across states, whereas for DyNeMo we retain the original values. We threshold the edges by selecting the top 2% of edges irrespective of sign.

#### Network response

Post-hoc, we perform an event-related response analysis on the inferred network activity (state/mode time courses for the HMM/DyNeMo). We calculate the *network response* by epoching and trial-averaging the state/mode time courses. For the HMM, the state time course corresponds to the Viterbi path (binarised state probability time course). For DyNeMo, the mode time course corresponds to the maximum a posteriori estimate for the mixing coefficient at each time point. Importantly, since each state/mode describes a network with specific oscillatory activity, a network response represents a set of underlying oscillatory responses that occur across that network. This is shown in Figures 3D and E. The result of a dynamic network analysis is a 1D time course for the network response for each subject and state/mode.

**Figure 3.**
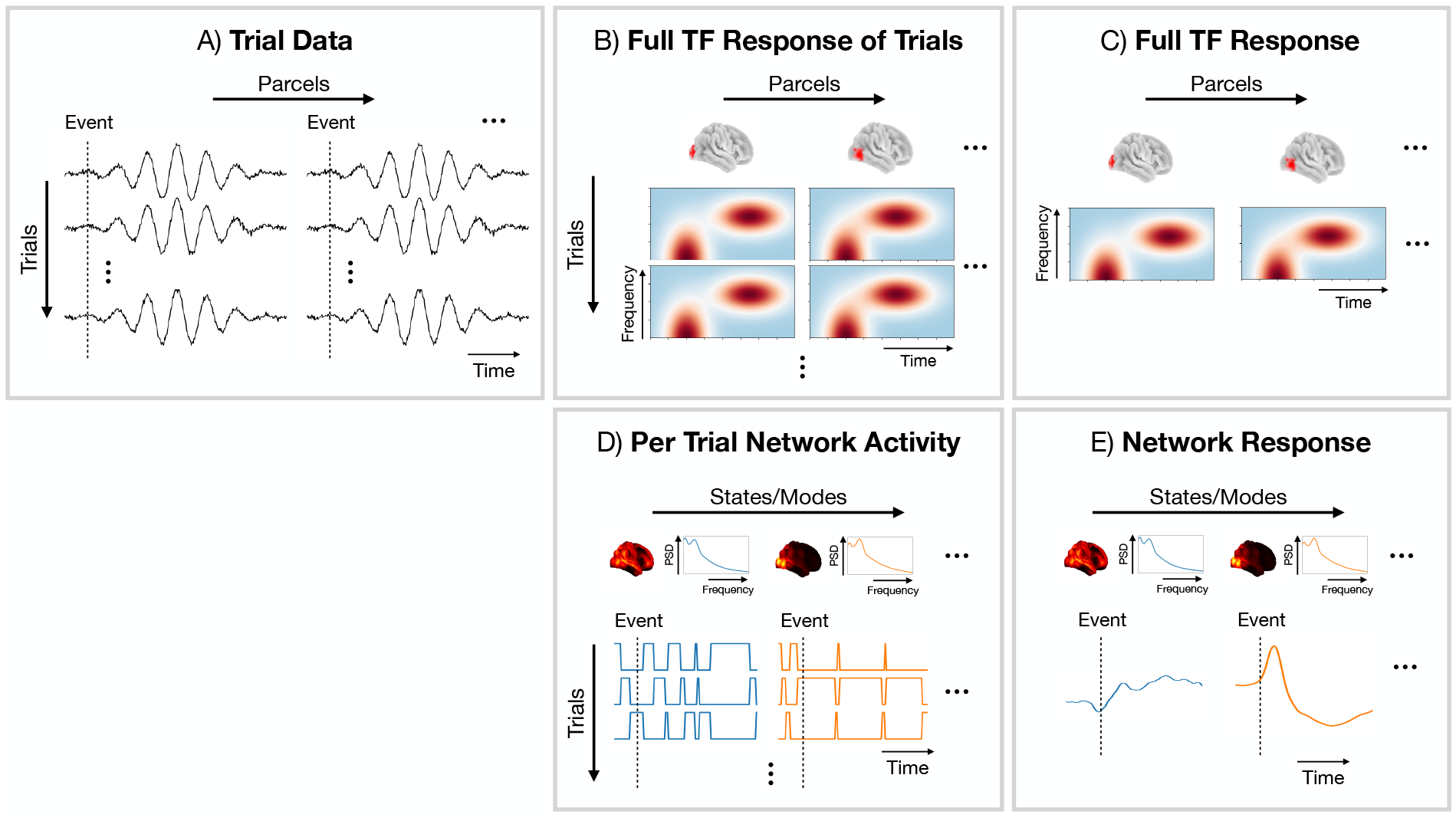
Comparison of a conventional parcel-based and network approach for oscillatory response analysis. A) Parcel time courses epoched around events of interest (trials). B) Full TF response of individual trials at the parcel level. C) Trial-averaged full TF response at the parcel level. D) Network activity (state/mode time course) epoched around events of interest. E) Network response (trial-averaged state/mode time course).

Similar to the TF response, we baseline correct the network response by subtracting the average value 100 ms before *t* = 0 s. We use the same approach as we did for the TF response to test for statistical significance (non-parametric permutation testing with a GLM). Note, the network response contains both the evoked and induced oscillatory response. The inability of the dynamic network models in distinguishing between evoked and induced oscillatory responses is a limitation of the network approach.

## 3 Results

We will first compare the dynamic network perspective provided by the HMM and DyNeMo using simulated data in Section 3.1. Following this, we will compare the different methods for analysing task data using the real MEG dataset (described in Section 2.1).

### 3.1 Simulation

To illustrate DyNeMo’s ability to identify overlapping network activations, we trained both dynamic networks models on the simulated data described in Section 2.1.1. The ground truth simulated power map and PSD for each network is shown in Figures 4A.II and A.III respectively. The corresponding ground truth parcel-level brain activity is shown in Figure 4A.I. The dashed vertical lines in Figure 4A.I show the onset of visual (green) and motor (blue) network activations.

**Figure 4.**
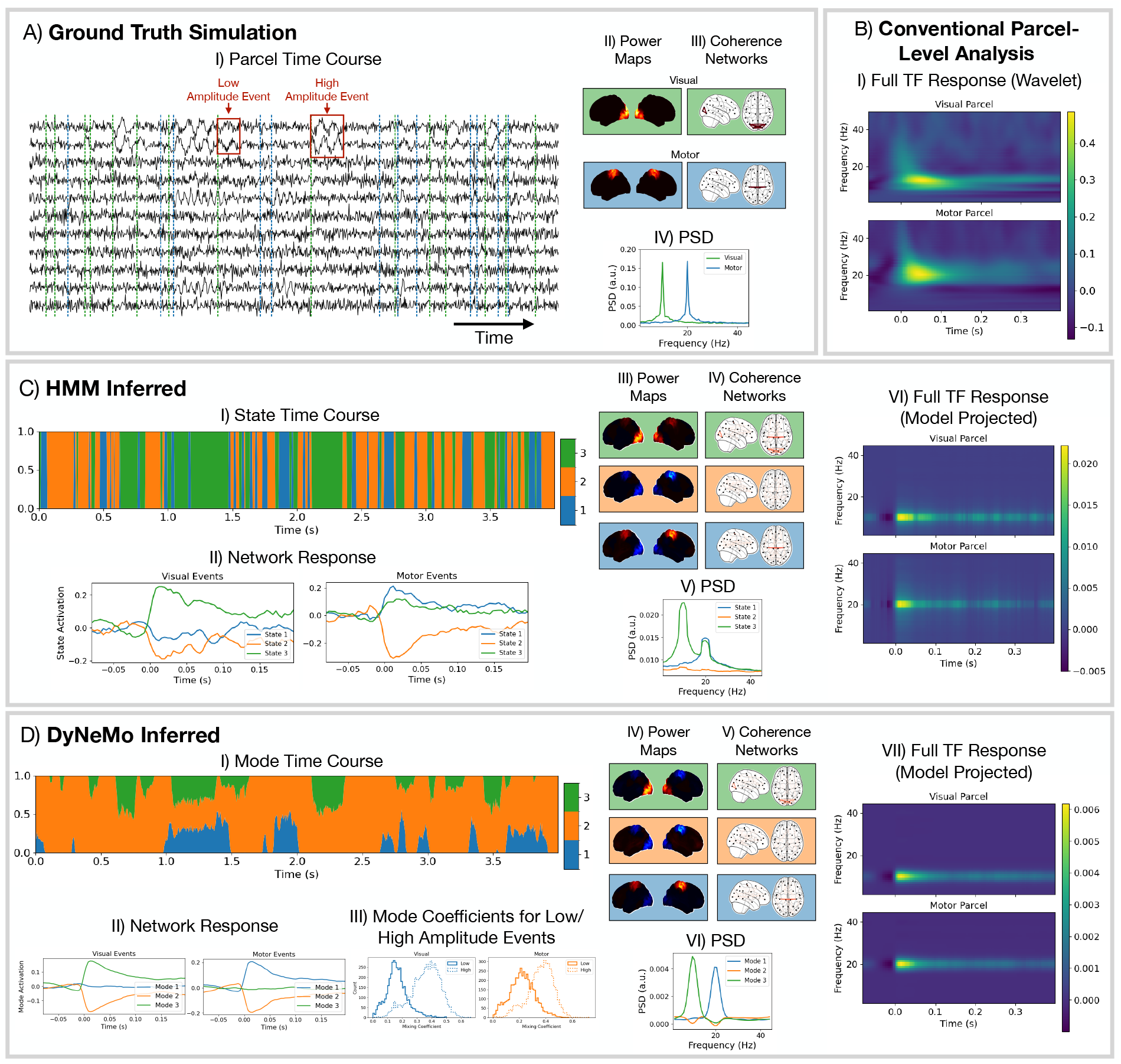
The assumption of mutual exclusivity harms the HMM’s ability to model the data when there are co-activating bursts of oscillatory activity. A) Ground truth simulation of parcel-level data: a subset of the parcel time courses (I) and properties of the networks (II-IV). B) Conventional parcel-level analysis applied to the simulated data. C) HMM network analysis applied to the simulated data. D) DyNeMo network analysis applied to the simulated data. C.VI) and D.VII) were calculated by multiplying the state/mode PSDs by the network response for the HMM and DyNeMo respectively.

Using the simulated parcel-level data (shown in Figure 4A), we performed a conventional TF response analysis selecting a parcel in the occipital cortex (Figure 4B.I, top) and motor cortex (B.I, bottom). We observe the conventional single-parcel full TF analysis correctly identifies the oscillatory frequency of each burst. Note, there is also a wide-band activation across many frequencies at the onset of the valid 10 (or 20) Hz power increase at approximately 100 ms, which is an artefact of the spectral estimation method (wavelet transform).

Next, we trained the HMM on the simulated parcel-level data. The perspective provided by the HMM is shown in Figure 4C. We can see from the network response (C.II) that there is a clear response to the onset of visual and motor bursts. However, we see that multiple states are activated in response to each event type. We also see from the power maps (C.III) that the activity from both event types is present in each state. Looking at the model projected full TF response (C.V), which is calculated by multiplying the network response time course by the PSD for each state and summing, we can see the HMM is still a good model for the underlying oscillatory dynamics, just its decomposition of activity into states prevents it from identifying the ground truth description of the data. Note, increasing the number of states in the HMM does not help it identify the ground truth networks in the simulation.

Finally, we trained DyNeMo on the simulated parcel-level data. The perspective provided by DyNeMo is shown in Figure 4D. We see from the network response (D.II) that DyNeMo is better able to isolate bursts of oscillatory activity into separate modes. We also see from the power maps (D.IV) that DyNeMo correctly identifies the simulated networks and the model projected full TF response (D.VI) is a good representation of the ground truth oscillatory content of the simulated data. In addition to the correct identification of event timings, DyNeMo is also able to learn the amplitude of each event type via the value of the mixing coefficients. Figure 4D.III shows the distribution of mode mixing coefficient values for when high and low-amplitude visual events occur (left) and motor events occur (right). We see when there is a low-amplitude event the mean of the mode mixing coefficient distribution is much lower than for high amplitude events. Note, using less modes than the number of networks in DyNeMo leads to the visual and motor networks combining into a single mode. Specifying more than three modes, DyNeMo learns only three modes are needed to describe the data and the extra modes have virtually no activation.

### 3.2 Real Task MEG Dataset

There are five unique events in this dataset: the presentation of a famous face, unfamiliar face, scrambled image, left button press and right button press (see Figure 1). We will study the average response (across trials, subjects and sessions) for different event types. Specifically, we will look at:

- What’s the response to all visual stimuli? (Section 3.2.1.)
- What’s the difference in response between faces and scrambled images? (Section 3.2.2.)
- What’s the difference in response between famous and unfamiliar faces? (Section 3.2.3.)
- What’s the response to a button press (left or right)? (Section 3.2.4.)

#### 3.2.1 Visual Stimuli: All

Turning to the real MEG dataset. First, we look at the average response over all visual stimuli.

##### Single-channel analysis reveals a fast oscillatory response to visual stimuli in the *α*-band in the occipital lobe and *θ*-band in the frontal lobe

Figure 5 shows the TF response calculated using conventional single-channel analysis. To be able to show such a parsimonious result, we have pre-selected a single sensor/parcel in the occipital lobe, as this is an area known to be involved with visual processing [23], and in the frontal lobe. At the sensor-level (Figure 5A), we can see a short-lived increase in *α*-band (8-12 Hz) activity followed by a decrease relative to baseline and a long-lived increase in *θ*-band (*<* 8 Hz) activity in the frontal lobe. We see a consistent description in source space (Figure 5B). Both analyses indicate a very fast response to the task, on the order of 100 ms, which is consistent with existing literature [24].

**Figure 5.**
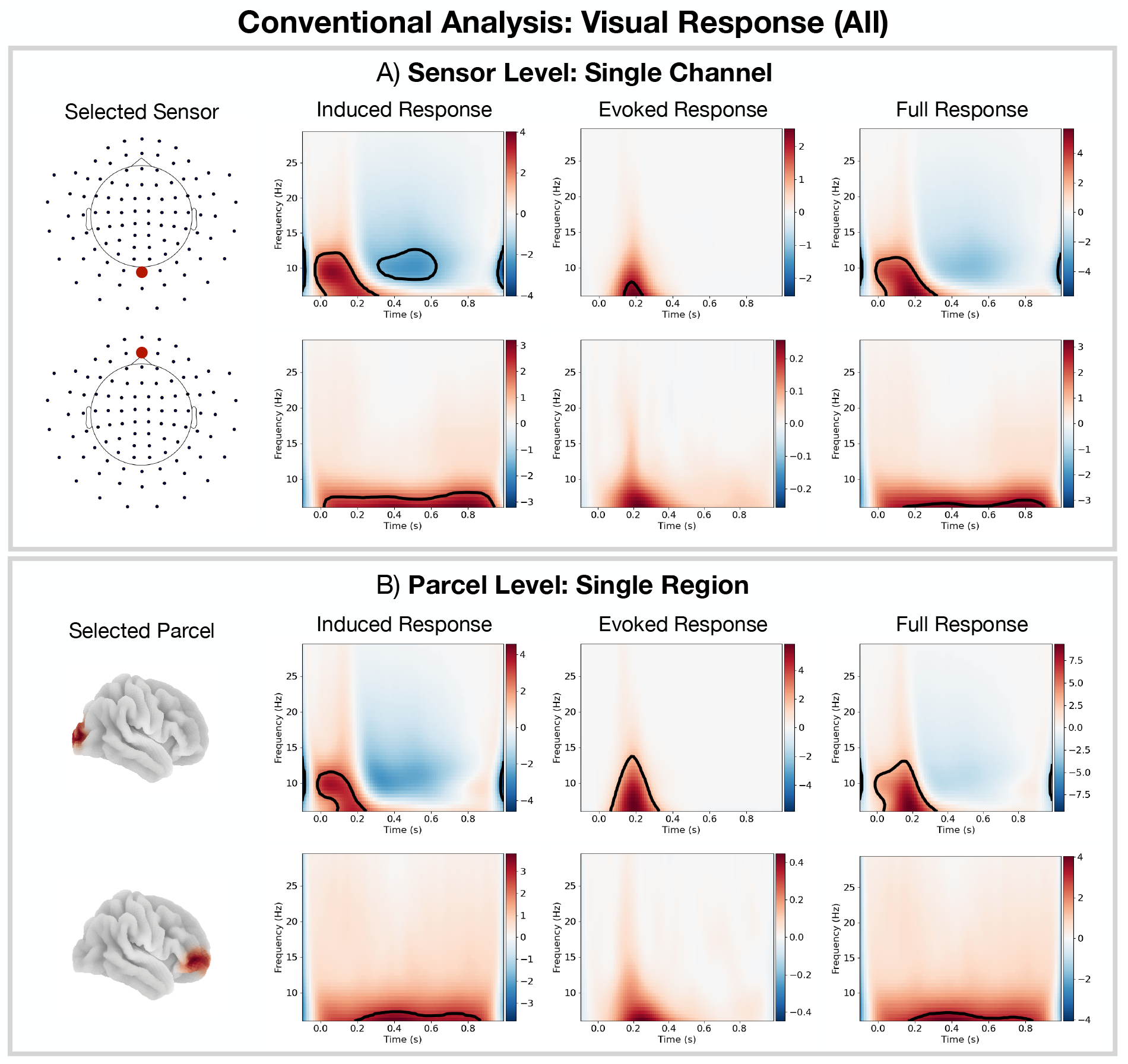
Conventional TF analysis reveals a fast oscillatory response to the task. For all visual stimuli: A) Conventional sensor-level analysis. B) Conventional parcel-level analysis. The black outline in TF plots indicate clusters with *p*-value *<* 0.05. Note, we had to pre-select the sensors/parcels of interest in this analysis.

##### Functional brain networks of oscillatory activity are identified by the HMM and DyNeMo directly from the data

Figure 6 shows the networks inferred by the HMM (A) and DyNeMo (B). Figure 6A.I and B.I show the spatial distribution of oscillatory power for each network as well as the phase-locking connectivity (coherence). We identify well known functional networks found in MEG data [18]. Of particular interest in this work is HMM states 2 and 5, which represent visual networks with *α*-band activity; state 3, which represents a frontotemporal network that is often associated with higher-order cognition [23] with *δ/θ*-band (1-8 Hz) activity; and state 4, which is a sensorimotor network with *β*-band (13-30 Hz) activity. HMM state 1 is identified as the default mode network with wide-band (1-45 Hz) activity [9]. The remaining state (6) represents a suppressed wide-band power network, which in previous HMM analyses has often been ignored. In this work, we will explore the relevance of this network in relation to the task and its connection to the DyNeMo networks. The DyNeMo networks that are of particular interest are: mode 2, which is a visual network with *α*-band activity; mode 3, which is a frontotemporal network with *δ/θ*-band activity; and modes 4 and 5, which respectively represent right and left lateralised sensorimotor networks with *β*-band activity. The remaining networks are mode 1, which is a left temporal network with *θ/α*-band (4-12 Hz) activity often associated with language [25] and mode 6, which is a suppressed power background network that does not show any additional oscillatory activity compared to the static (time-averaged) PSD [11]. Figure 6C shows the average values for the (renormalised) DyNeMo mode time course when each HMM state is active. This signifies the amount each DyNeMo mode contributes to an HMM state. We see multiple DyNeMo modes combine to form an HMM state. Note, the networks presented in Figure 6 can be reliably found in this dataset, see SI Figure S5.

**Figure 6.**
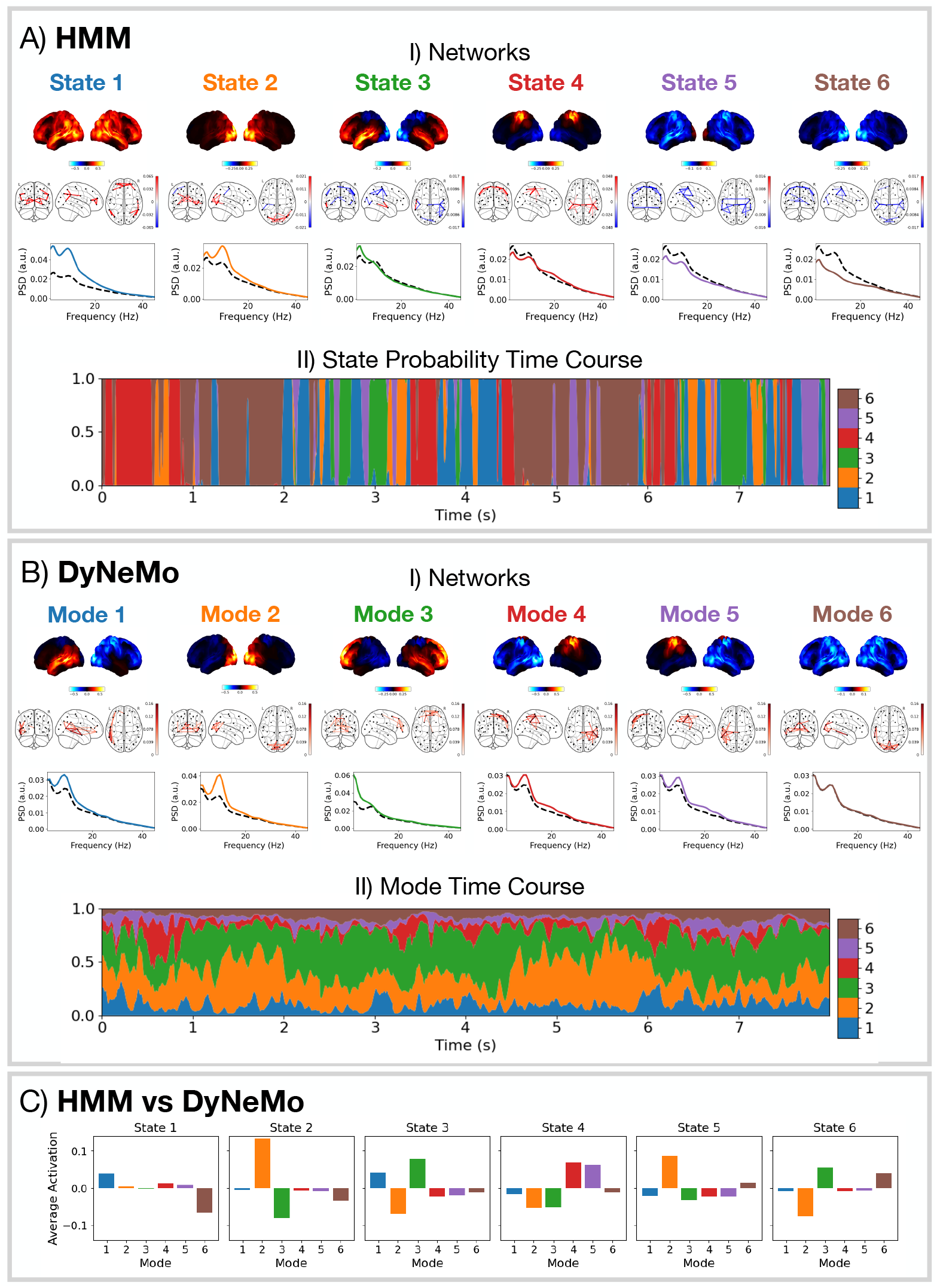
DyNeMo and the HMM provide a fundamentally different network perspective. A.I) Network states inferred by the HMM: power maps (top), coherence networks (middle), and PSDs (solid coloured line) with the static PSD (dashed black line) (bottom). A.II) State probabilities for the first 8 seconds for the first session. The width of each colour represents the probability of a particular state. B.I) Network modes inferred by DyNeMo: power maps (top), coherence networks (middle), and PSDs (bottom). B.II) Renormalised mode time course (weighted by trace of each mode covariance [11]) for the first 8 seconds for the first session. C) Time-average DyNeMo renormalised mode time course when each HMM state is active minus the average across all time points.

##### Network responses to task are identified with millisecond temporal resolution

Figure 7A.II and B.II show network responses in the form of trial-averaged epoched state/mode time courses. Importantly, since each state/mode describes a network with specific oscillatory activity, a network response represents a set of underlying oscillatory responses that occur across that network. Both the HMM and DyNeMo were able to identify networks with millisecond-level dynamics directly from the data. DyNeMo provides a much simpler network response compared to the HMM. We see an initial visual network activation, followed by a frontotemporal network activation combined with a visual network deactivation. The reason we see many significant network activations and deactivations for the HMM is likely due to the mutual exclusivity, where for a network to activate, all other networks must deactivate. Lifting this constraint leads to the more parsimonious description provided by DyNeMo. Note, DyNeMo’s network response looks smoother than the HMM because DyNeMo has the ability to model both the amplitude and timing of network activations via the mode time course (see Figure 4D.III), whereas the HMM can only model the timing with a binary state time course. Comparing the network responses (Figure 7A.II and B.II) with a conventional single-parcel analysis (full TF response in Figure 5B), we see oscillatory activity in the occipital lobe parcel is actually part of a wider visual network. Using the HMM/DyNeMo network description we can project a model estimate for the oscillatory activity at a single parcel^6^. This is shown in SI Figure S5. We can see both the HMM and DyNeMo provide a good description of the oscillatory response to visual stimuli for a single parcel.

**Figure 7.**
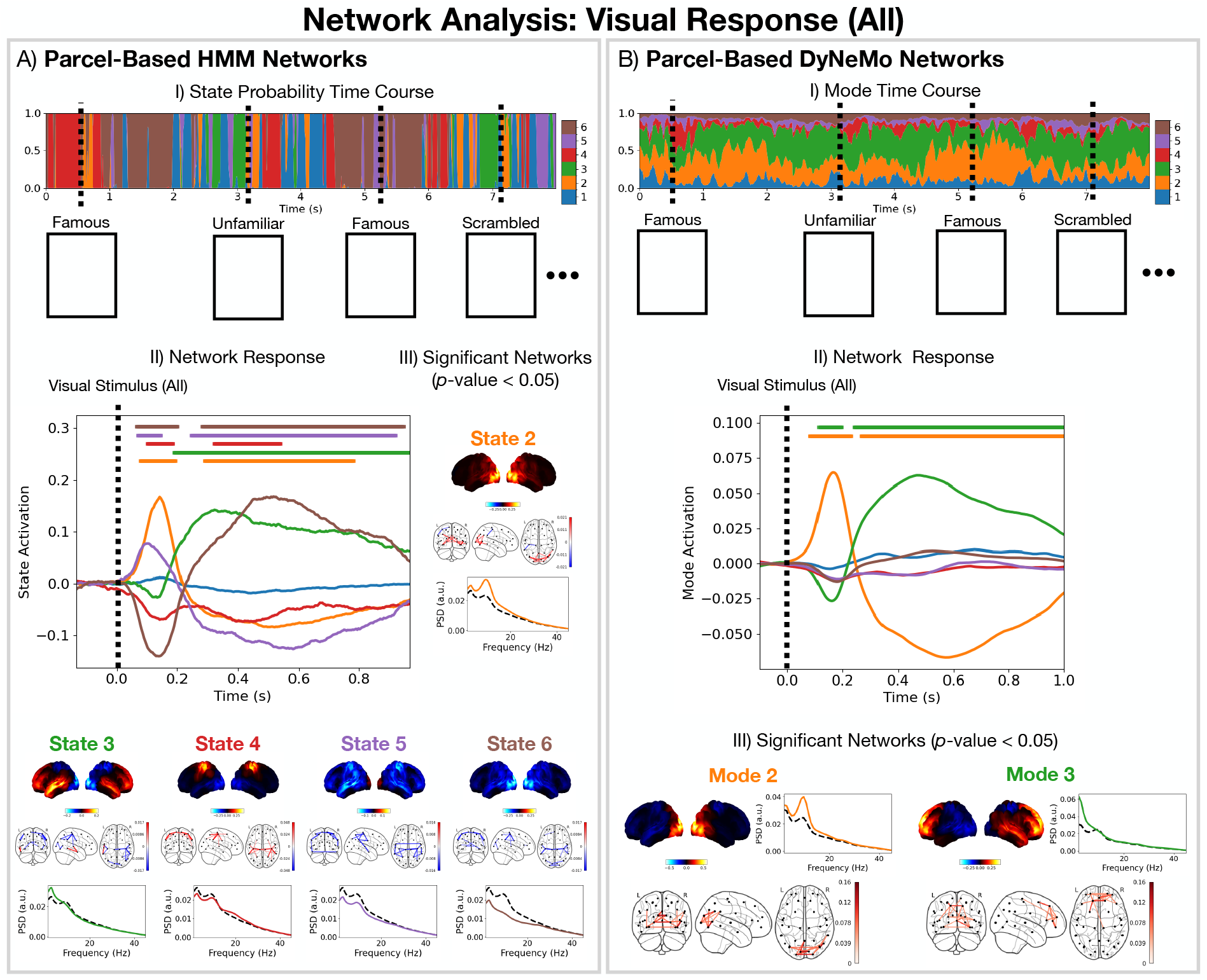
Both the HMM and DyNeMo reveal a millisecond resolution dynamic network-level perspective of brain activity in response to the task, with DyNeMo providing a more parsimonious decomposition. For all visual stimuli: A) Parcel-based HMM analysis: (I) The inferred state probabilities for the first 8 seconds of the first session. The dashed vertical lines indicate when a particular visual stimulus was presented; (II) Network response. The colour of each line indicates its state; A.III) Properties of the networks that show a significant response (*p*-value *<* 0.05). B) Parcel-based DyNeMo analysis: (I) The inferred (renormalised) mode time course; (II) Network response; (III) Properties of the networks that show a significant response (*p*-value *<* 0.05). The horizontal bars above each time course in (A.II) and (B.II) indicate the time points with *p*-value *<* 0.05.

#### 3.2.2 Visual Stimuli: Faces vs Scrambled

Next, we examine a more subtle task effect by looking at the difference in response between the presentation of a face vs scrambled image (the response to faces (famous and unfamiliar) minus the response to scrambled images). Studying this process allows us to understand the cognitive processes that occur in the detection and identification of a face in contrast to non-facial images.

##### Dynamic network approaches (HMM and DyNeMo) can identify a late network response that conventional methods fail to detect

Figures 8A and B show the difference in TF response for faces vs scrambled calculated using conventional methods at the sensor and parcel level respectively. We see using conventional methods that we fail to identify any significant induced TF responses in posterior sensors/parcels (Figure 8A and B). We do observe a significant increase in evoked and full TF response but only at the parcel level (Figure 8B). SI Figure S7 shows we fail to observe any significant TF responses in frontal sensors/regions. Turning to the dynamic network models, which describe the full TF response to the task, we see both the HMM and DyNeMo show a larger activation of the visual network for faces over scrambled images (Figure 8C and D, right). Additionally, the HMM is able to identify a late activation of a suppressed power network (state 6, Figure 8C, right). Figure 8D (middle) shows the time-averaged value for each DyNeMo (renormalised) mode time course for time points in the data when HMM state 6 is active. We see HMM state 6 can be represented as a mixture of DyNeMo modes 2, 3 and 6 and that the observed late response seen with the HMM (state 6 in Figure 8C, right) is the combination of a visual network (mode 2) deactivation and frontotemporal network (mode 3) activation in DyNeMo (Figure 8D, right).

**Figure 8.**
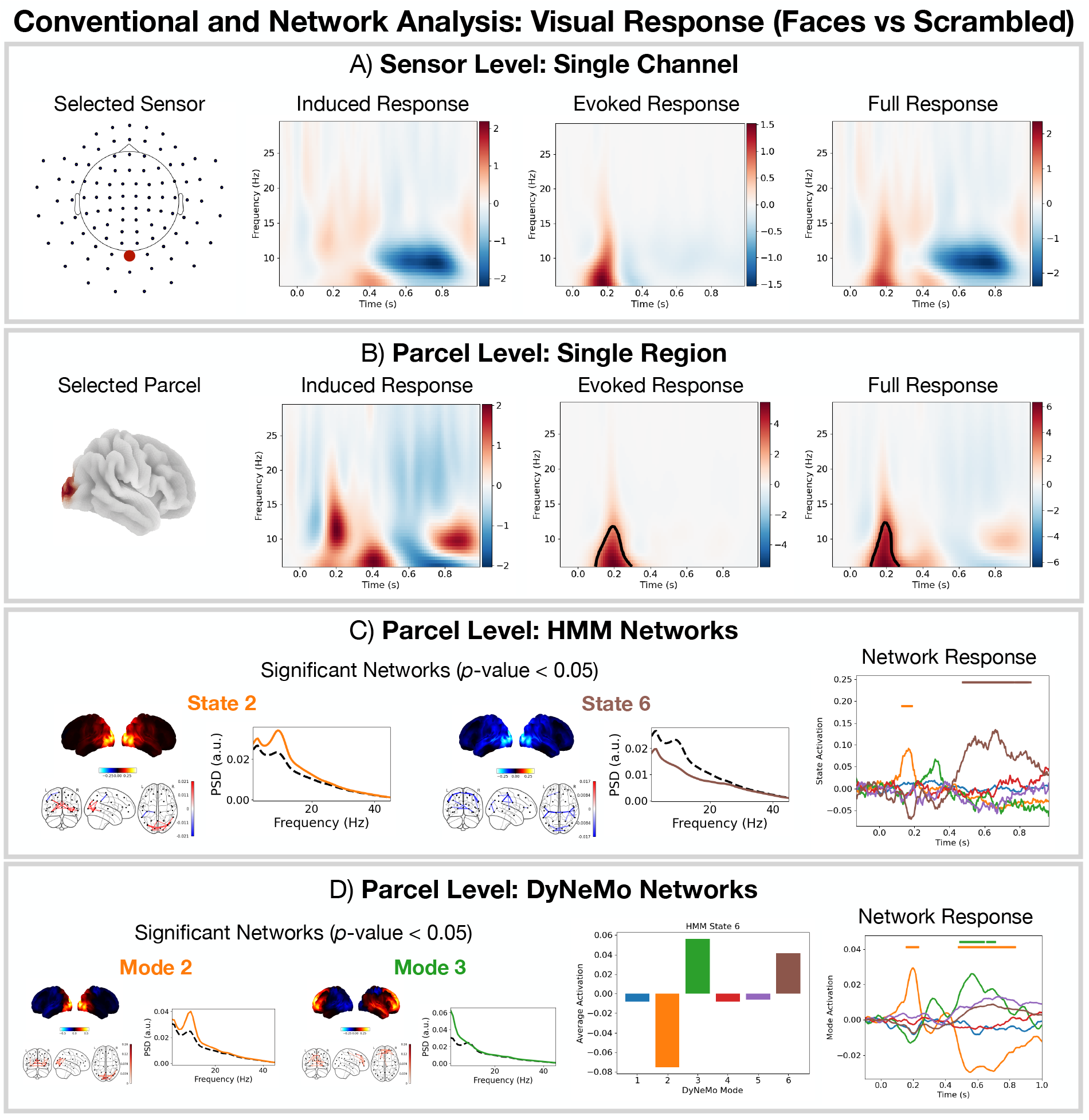
Unique combinations of DyNeMo modes are modelled as distinct states in the HMM. For faces (famous and unfamiliar) vs scrambled images: A) Conventional single-channel sensor-level analysis. B) Conventional single-region parcel-level analysis. C) Parcel-based HMM network analysis. D) Parcel-based DyNeMo network analysis. The black outline in the TF plots in (A) and (B) indicate clusters with *p*-value *<* 0.05. The horizontal bars above each time course in (C) and (D) indicate the time points with *p*-value *<* 0.05. The bar chart in (D) shows the activation of each DyNeMo mode when HMM state 6 is active.

##### Dynamic network analyses (HMM and DyNeMo) reveal a sequence of oscillatory network activations related to the cognitive processing of faces

Studying the difference between the network response to the presentation of a face (famous and unfamiliar) and scrambled images allows us to understand the network activations that occur during the processing of faces. First turning our attention to the network description provided by the HMM (Figure 8C, right), we see there is an early visual network response (state 2, occipital *α*-band activity) that lasts approximately 100 ms, which could represent a bottom-up response to the detection of a face. Following this, we see an activation of a frontotemporal network (state 3, frontotemporal *δ*/*θ*-band activity) that lasts approximately 100 ms. Following this, there is a long-lived (300-400 ms) activation of the suppressed visual network (state 6, suppressed occipital *α*-band activity). The DyNeMo perspective (Figure 8D, right), shows something similar, however, it is able to identify that the late suppression of the visual network (deactivation of mode 2) occurs simultaneously with an activation of a frontotemporal network (mode 3). This provides a new insight that the suppressed activity in the occipital cortex maybe linked to the activity in frontotemporal regions. Such activity could represent a top-down feedback response that occurs in the identification of faces. This description is consistent with existing literature on face recognition in the primate visual system [13].

#### 3.2.3 Visual Stimuli: Famous vs Unfamiliar

Next, we look at an even more subtle effect by comparing the difference in response between famous and unfamiliar faces (the response to famous faces minus the response to unfamiliar faces).

##### Only DyNeMo reveals a deactivation of the visual network and activation of the frontotemporal network for famous vs unfamiliar faces

Figure 9A and B show the results of a conventional TF analysis at the sensor and parcel-level respectively for the famous vs unfamiliar contrast. We see no significant TF response. Looking at the HMM analysis for this contrast in Figure 9C, we see a hint that there may be a response in the suppressed power network (state 6). However, this response is unconvincing. Comparing this to the network response identified by DyNeMo in Figure 9D, we see a much clearer response with a deactivation of the visual network (mode 2) and activation of the frontotemporal network (mode 3).

**Figure 9.**
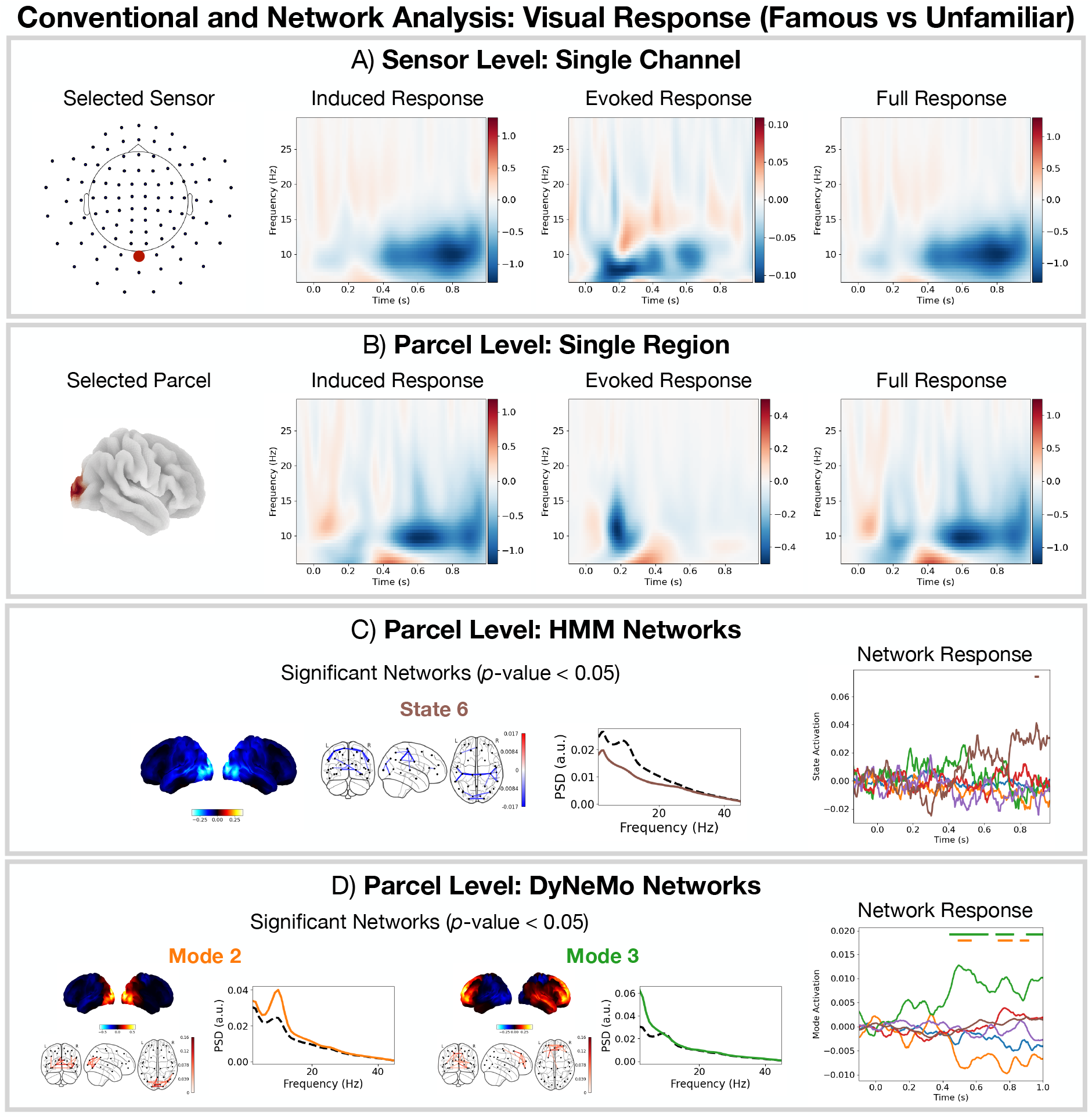
Modelling the data using DyNeMo can reveal subtle differences in brain activity missed by conventional methods and the HMM. For famous vs unfamiliar faces: A) Conventional single-channel sensor-level analysis. B) Conventional single-region parcel-level analysis. No significant time points or frequencies were found with conventional analysis. C) Parcel-based HMM network analysis. D) Parcel-based DyNeMo network analysis. The horizontal bars above each time course in (C) and (D) indicate the time points with *p*-value *<* 0.05.

##### The identification of faces (famous vs unfamiliar) occurs in the late network response

Focusing on the network response provided by DyNeMo (Figure 9D) we do not see the early visual network (mode 2) activation that we saw in the faces vs scrambled contrast (Figure 8D). This suggests the detection of a face occurs early on (around 100 ms after the presentation of the image). Following this, the network response shown in Figure 9D suggests the identification of the face (whether is it a famous or unfamiliar face) occurs after 400 ms and involves the frontotemporal network (mode 3) and visual network (mode 2). This description is consistent with existing literature [13].

#### 3.2.4 Button Press: Left vs Right

The final contrast we study is the response to a button press. Note, the experiment involves two types of button presses: a left button press executed with the left index finger and right button press executed with the right index finger.

##### DyNeMo identifies unilateral components of the sensorimotor network, providing a better model of the data than the HMM

Figure 10A and B respectively show the results of a conventional single-channel (sensor/parcel) analysis. Here, we epoch around all button presses (left or right). We selected a sensor/parcel near the motor cortex as we expect this area to be involved in executing a button press [23]. We observe the expected post-movement *β*-rebound a few hundred milliseconds following the button press [26]. We see a consistent network description with both the HMM (Figure 10C) and DyNeMo (Figure 10D). Note, the activation of the visual networks after the button press is due to the start of the next trial. An interesting feature of the network description provided by DyNeMo is that the sensorimotor network has been split into two unilateral components (modes 4 and 5). Epoching the mode time courses for these two modes around left and right button presses, which were executed with left and right index fingers respectively (Figure 10E), we see the contralateral network activates for each button press. This is in contrast to the HMM, where a single bilateral sensorimotor network activates in response to both left and right button presses.

**Figure 10.**
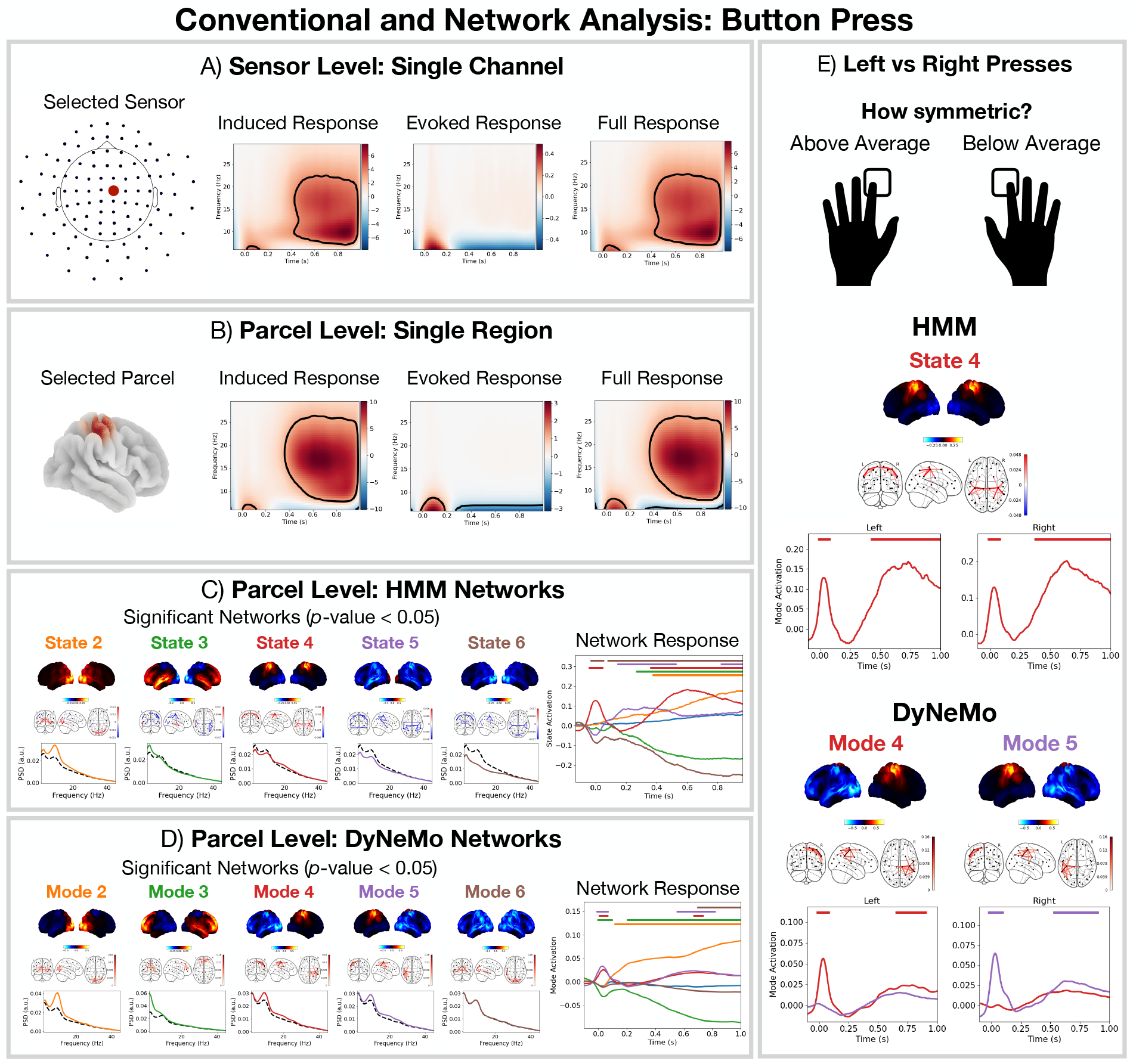
DyNeMo offers a more precise dynamic network description, inferring significant unilateral sensorimotor networks that are missing from the HMM. For the button press events: A) Conventional single-channel sensor-level analysis. B) Conventional single-region parcel-level analysis. C) Parcel-based HMM networks analysis. D) Parcel-based DyNeMo network analysis. E) Network responses for the HMM/DyNeMo epoched around left and right button presses. The black outline in the TF plots in (A) and (B) indicate clusters with *p*-value *<* 0.05. The horizontal bars above each time course in (C), (D) and (E) indicate the time points with *p*-value *<* 0.05.

## 4 Discussion

### Sensor vs source-space analysis

In Figures 5, 8-10 we compared a conventional single-channel TF analysis in sensor and source (parcel)-space. We observed both methods provided consistent descriptions of the timing and oscillatory content of the response to the task. The main advantage of performing the analysis in source space is the ability to isolate the response in a particular region in the brain, which can aid with interpretability. For example, in Figure 5B we were able to show the oscillatory response to visual stimuli occurs in the occipital lobe. An additional benefit of performing analysis in source-space is that the process of source reconstruction incorporates information regarding the head position relative to the sensors, which could provide a useful denoising effect.

### Disadvantages of a conventional analysis

The conventional event-related oscillatory response analysis of task data uses region-by-region TF decompositions. It is simple, quick and easy to compute. However, it is limited in what it can achieve. It only looks at a single sensor or region of interest at a time. Additionally, there can be a large number of sensors/parcels, which means we can end up performing a large number of multiple comparisons when we test for statistical significance, when we cannot sensibly pre-specify a small set of sensors/parcels of interest. This can drastically reduce our sensitivity to subtle effects. For example, we could not identify any differences in the response to famous vs unfamiliar faces in Figure 9A and B.

### Advantages of a dynamic network analysis

Adopting a dynamic network approach can provide a useful perspective on how the brain responses to a task. Rather than doing many parcel-based TF analyses to oscillatory responses, we can instead identify how large-scale networks respond to the task. Since each state (with the HMM) or mode (with DyNeMo) describes a network with specific oscillatory activity, these network responses represent a set of underlying oscillatory responses that occur across that network that we can interrogate. E.g., with a dynamic network approach we can understand which large-scale networks underpin single-region oscillatory changes. Also, once we know which networks activate or deactivate in response to a task, we can look up existing literature relating to the network and understand its function in a wider context. In Figure 7, we identified a visual and frontotemporal network is involved in the task we studied.

Another useful advantage is there’s no need to specify the oscillatory frequencies of interest. Both the HMM and DyNeMo can learn networks with distinct oscillatory activity directly from the data. We also show in Figure 7 that we can resolve dynamics with the same resolution as event-related TF analyses, meaning we do not sacrifice any temporal precision with the dynamic network perspective. Note, the network response reflects both the evoked and induced TF response.

Decomposing brain activity into a low number of states (with the HMM) or modes (with DyNeMo) acts as a dimensionality reduction technique, which can aid with interpretation. In this work, we chose 6 to give us a low-dimensional network description. This also allows us to reduce the number of multiple comparisons we perform, which can increase our sensitivity to subtle effects. For example, only with the network models did we see a difference in the response between faces and scrambled images (Figure 8).

### The dynamic networks identified by the HMM and DyNeMo can be reliably inferred

As with many modern machine learning models, the HMM and DyNeMo can suffer from a local optima issue, where slightly different networks can be inferred from the same data. With good initialisation, improved training techniques and larger datasets, this has been minimised [18]. Empirically, we find selecting the best run^7^ (the one with the lowest loss^8^) from a set of 10 runs robustly finds a reproducible network descriptions of the data. We also find group-level analysis, such as the network responses, are robust. In Figure S5, we show the same analysis repeated on 3 sets of 10 runs and demonstrate the same networks and conclusions are obtained with each.

The DyNeMo networks (Figure 6B) appear to be more localised compared to the HMM networks (Figure 6A). This raises the question whether the localised connectivity patterns we observe are due to volume conduction. The measure for connectivity that we use is coherence, which quantifies the level of phase stability between two oscillations irrespective of the value for the phase difference. Volume conduction results in zero phase lag activity, which we remove in the symmetric orthogonalisation step following parcellation. Therefore, we conclude the coherence we observe is not due to volume conduction.

### DyNeMo provided the most parsimonious network description of the methods tested

Both the HMM and DyNeMo are state-of-the-art methods for identifying dynamic networks in neuroimaging data. However, we showed in simulation (Figure 4) that the assumption of mutually exclusive states may compromise the network description provided by the HMM. DyNeMo overcomes this limitation by allowing networks to overlap at a given time point. The network description provided by DyNeMo appears to be much more straightforward to interpret. We saw in Figure 7B, that the network response to visual stimuli was isolated to just a couple networks (modes 2 and 3) with localised power activations. Whereas with the HMM (Figure 7A) many networks (states 2-6) show a significant response to the task. This is likely an artefact of the assumption of mutual exclusivity, which for a state to activate requires all other states to deactivate.

### DyNeMo offered the most precise network descriptions of the methods tested

In this task, participants perform a button press with either their left or right index finger. When we compared the network response to left and right button presses separately, we saw DyNeMo was able to identify unilateral sensorimotor networks contralateral to movements (modes 4 and 5) whereas the HMM did not (Figure 10E). This network description was learnt directly from the data, without any information regarding the button press being fed into model. Such a description could be useful for studying lateralisation of function [28] or inter-hemispheric connectivity [29].

### Alternative approaches

DyNeMo is not the only approach that can model overlapping spatial patterns. For example, Hyvärinen et al. proposed ‘Fourier-ICA’, which applies ICA to the stacked short-time Fourier transform of each channel in M/EEG data [30]. This approach is able to find overlapping patterns of time-frequency modes across channels. However, it does not model the interaction between channels or brain regions. DyNeMo in contrast is an explicit model for the interaction between channels or brain regions; in the latter case providing a task response in terms of functional connectivity or graphical network dynamics. This is achieved via the cross-spectral density modelling in the covariance matrix of time-delay embedded data, and DyNeMo is free to use this information to help infer the best spatiotemporal modes that describe the data. DyNeMo also benefits from an explicit temporal model (the ‘model RNN’ [11]), which helps in inferring the mode time courses, whereas the sources in Fourier-ICA (analogous to the mode time courses in DyNeMo) are determined only by the maximisation of non-Gaussianity. In both approaches, choices need to be made in preparing the training data. For Fourier-ICA, it is the parameters used to calculate the short-time Fourier transform. For DyNeMo, it is the choice of time-delay embedding and PCA parameters. Another more recent approach was proposed by Power et al. in [31], which used convolutional dictionary learning on MEG data to identify repeated ‘atoms’ of temporally overlapping spatial patterns across channels in response to a task. One advantage of this approach is its ability to learn the waveform shape of the response in the raw training time series, which is not possible with Fourier-ICA or DyNeMo. However, unlike DyNeMo this approach does not model the interaction between channels or brain regions.

## 5 Conclusions

Conventional event-related methods can be a useful technique for understanding the oscillatory response at a single region. However, a potentially more powerful technique that can give a whole brain perspective is a dynamic network approach. We have shown that two state-of-the-art methods: the HMM and DyNeMo can learn frequency-specific large-scale functional networks that activate in response to a task with millisecond resolution. Comparing the HMM with DyNeMo, we found DyNeMo provides a particularly useful network description for studying task data. We provide the scripts for reproducing the analysis presented in this report here: github.com/OHBA-analysis/Gohil2024_NetworkAnalysisOfTaskData.

## 6 Ethics Statement

The original study that collected the data [19] was approved by Cambridge University Psychological Ethics Committee. Written informed consent was obtained from each participant prior to and following each phase of the experiment. Participants also gave separate written consent for their anonymised data to be freely available on the internet.

## 7 Data and Code Availability Statement

The data is publicly available [19]. Code to reproduce analysis presented in this manuscript is available here: https://github.com/OHBA-analysis/Gohil2024_NetworkAnalysisOfTaskData.

## 8 Author Contributions

CG: Conceptualisation; Methodology; Software; Validation; Formal Analysis; Data Curation; Writing - Original Draft; Writing - Reviewing & Editing; Visualisation. OK: Conceptualisation; Writing - Reviewing & Editing. RH: Software. MWJE: Conceptualisation. OPJ: Writing - Review & Editing; LTH: Writing - Review & Editing. AJQ: Conceptualisation; Methodology. MWW: Conceptualisation; Methodology; Data Curation; Supervision; Writing - Original Draft; Writing - Reviewing.

## 9 Declaration of Competing Interests

No competing interests.

## Footnotes

1 We often refer to the data collected during a task as *task data*.

2 This number of lags is sufficient in this simulation example because of the relatively low and narrowband oscillatory frequencies that are contained in the data. In real data, we typically use *±*7 lags because there are many more and boarder oscillatory frequencies present.

3 We also simulate the scenario of increased oscillatory power without the phase coupling by narrowband filtering white noise, see SI Figure S4 for this.

4 In the accompanying Python scripts, each session is referred to as a ‘run’.

5 Note, in this work we will reweight and renormalise the inferred mode coefficients using the trace of each mode covariance. This ensures the (renormalised) mode time courses account for the magnitude of each covariance. See [11] for further details.

6 This is done by multiplying the network response by the state/mode PSD.

7 A *run* is a single model trained from scratch.

8 In our case the loss is the *variational free energy* [27] for both DyNeMo the the HMM.

## Notes

https://github.com/OHBA-analysis/Gohil2023_NetworkAnalysisOfTaskData

### Competing Interest Statement

The authors have declared no competing interest.

### Summary of Updates

Simulation results and link to public repository containing example scripts was updated.

## References

[1] Buzsáki, G. (2006). Rhythms of the Brain. Oxford university press.

[2] Proudfoot, M., Woolrich, M. W., Nobre, A. C., & Turner, M. R. (2014). Magnetoencephalography. Practical neurology, 14 (5), 336–343.

[3] Luck, S. J. (2014). An introduction to the event-related potential technique. MIT press.

[4] Urai, A. E., Doiron, B., Leifer, A. M., & Churchland, A. K. (2022). Large-scale neural recordings call for new insights to link brain and behavior. Nature neuroscience, 25 (1), 11–19.

[5] Lin, A., Witvliet, D., Hernandez-Nunez, L., Linderman, S. W., Samuel, A. D., & Venkatachalam, V. (2022). Imaging whole-brain activity to understand behaviour. Nature Reviews Physics, 4 (5), 292–305.

[6] Bressler, S. L., & Menon, V. (2010). Large-scale brain networks in cognition: emerging methods and principles. Trends in cognitive sciences, 14 (6), 277–290.

[7] Fries, P. (2015). Rhythms for cognition: communication through coherence. Neuron, 88 (1), 220–235.

[8] Quinn, A. J., Vidaurre, D., Abeysuriya, R., Becker, R., & Woolrich, M. W. (2018). Task-evoked dynamic network analysis through hidden Markov modeling. Frontiers in neuroscience, 12, 349904.

[9] Vidaurre, D., Hunt, L. T., Quinn, A. J., Hunt, B. A., Brookes, M. J., Nobre, A. C., & Woolrich, M. W. (2018). Spontaneous cortical activity transiently organises into frequency specific phase-coupling networks. Nature communications, 9 (1), 2987.

[10] Rabiner, L., & Juang, B. (1986). An introduction to hidden Markov models. ieee assp magazine, 3 (1), 4–16.

[11] Gohil, C., Roberts, E., Timms, R., Skates, A., Higgins, C., Quinn, A., … & Woolrich, M. (2022). Mixtures of large-scale dynamic functional brain network modes. NeuroImage, 263, 119595.

[12] Colclough, G. L., Brookes, M. J., Smith, S. M., & Woolrich, M. W. (2015). A symmetric multivariate leakage correction for MEG connectomes. Neuroimage, 117, 439–448.

[13] Moeller, S., Freiwald, W. A., & Tsao, D. Y. (2008). Patches with links: a unified system for processing faces in the macaque temporal lobe. Science, 320 (5881), 1355–1359.

[14] Tallon-Baudry, C., & Bertrand, O. (1999). Oscillatory gamma activity in humans and its role in object representation. Trends in cognitive sciences, 3 (4), 151–162.

[15] Winkler, A. M., Ridgway, G. R., Webster, M. A., Smith, S. M., & Nichols, T. E. (2014). Permutation inference for the general linear model. Neuroimage, 92, 381–397.

[16] Vidaurre, D., Quinn, A. J., Baker, A. P., Dupret, D., Tejero-Cantero, A., & Woolrich, M. W. (2016). Spectrally resolved fast transient brain states in electrophysiological data. Neuroimage, 126, 81–95.

[17] Vidaurre, D., Abeysuriya, R., Becker, R., Quinn, A. J., Alfaro-Almagro, F., Smith, S. M., & Woolrich, M. W. (2018). Discovering dynamic brain networks from big data in rest and task. NeuroImage, 180, 646–656.

[18] Gohil, C., Huang, R., Roberts, E., van Es, M. W., Quinn, A. J., Vidaurre, D., & Woolrich, M. W. (2024). osl-dynamics, a toolbox for modeling fast dynamic brain activity. Elife, 12, RP91949.

[19] Wakeman, D. G., & Henson, R. N. (2015). A multi-subject, multi-modal human neuroimaging dataset. Scientific data, 2 (1), 1–10.

[20] Quinn, A.J., van Es, M.W.J., Gohil, C., & Woolrich, M.W. OHBA Software Library in Python (OSL) (0.1.1). Zenodo (2022). 10.5281/zenodo.6875060.

[21] Hillebrand, Hillebrand A., & Barnes, G. R. (2005). Beamformer analysis of MEG data. International review of neurobiology, 68, 149–171.

[22] Westner, B. U., Dalal, S. S., Gramfort, A., Litvak, V., Mosher, J. C., Oostenveld, R., & Schoffelen, J. M. (2022). A unified view on beamformers for M/EEG source reconstruction. Neuroimage, 246, 118789.

[23] Baars, B., & Gage, N. M. (2013). Fundamentals of cognitive neuroscience: a beginner’s guide. Academic Press.

[24] Carlson, T., Tovar, D. A., Alink, A., & Kriegeskorte, N. (2013). Representational dynamics of object vision: the first 1000 ms. Journal of vision, 13 (10), 1–1.

[25] Fedorenko, E., & Thompson-Schill, S. L. (2014). Reworking the language network. Trends in cognitive sciences, 18 (3), 120–126.

[26] Cheyne, D. O. (2013). MEG studies of sensorimotor rhythms: a review. Experimental neurology, 245, 27–39.

[27] Bishop, C. M., & Nasrabadi, N. M. (2006). Pattern recognition and machine learning (Vol. 4, No. 4, p. 738). New York: springer.

[28] Mutha, P. K., Haaland, K. Y., & Sainburg, R. L. (2012). The effects of brain lateralization on motor control and adaptation. Journal of motor behavior, 44 (6), 455–469.

[29] Krupnik, R., Yovel, Y., & Assaf, Y. (2021). Inner hemispheric and interhemispheric connectivity balance in the human brain. Journal of Neuroscience, 41 (40), 8351–8361.

[30] Hyvärinen, A., Ramkumar, P., Parkkonen, L., & Hari, R. (2010). Independent component analysis of short-time Fourier transforms for spontaneous EEG/MEG analysis. NeuroImage, 49 (1), 257–271.

[31] Power, L., Allain, C., Moreau, T., Gramfort, A., & Bardouille, T. (2023). Using convolutional dictionary learning to detect task-related neuromagnetic transients and ageing trends in a large open-access dataset. NeuroImage, 267, 119809.

