## Supplementary information for "Dynamic Network Analysis of Electrophysiological Task Data"

### 1 Supplementary Information (SI)

#### 1.1 Statistical Significance Testing with a GLM

In this work, we use non-parametric permutations with a GLM (General Linear Model) to test for statistical significance. We fit the model:

$$\boldsymbol{y} = \boldsymbol{X}\boldsymbol{\beta} + \boldsymbol{\epsilon}, \tag{1}$$

where $\boldsymbol{y}$ [subjects $\times$ features] is the target data, $\boldsymbol{X}$ [subjects $\times$ regressors] are regressors, $\boldsymbol{\beta}$ [regressors $\times$ features] are regression coefficients and $\boldsymbol{\epsilon}$ [subjects $\times$ features] are the residuals. The features can be any subject-specific quantity of interest. In this report, the features are the response (evoked/induced TF response or state/mode activation) at each time point for each channel (sensor, parcel, state or mode).

We fit one regressor with a constant value of 1 for all subjects. This results in the regression coefficients giving the mean across subjects. Note, when fitting the GLM we treat each run as an individual subject. To test if the observed group mean is significantly different to zero, we look at the following COPE (Contrast of Parameter Estimate):

$$|\boldsymbol{\beta}| > 0, \tag{2}$$

where $|.|$ denotes the absolute value. We do this by applying 'sign-flip' permutations to the regressor. Each entry in the regressor has a 50% chance of being multiplied by -1. We perform this permutation 1,000 times and record the maximum COPE ($\max(|\boldsymbol{\beta}|)$) - this accounts for multiple comparisons. The 1,000 values for the maximum COPE provide a null distribution under the hypothesis that there is no response (equal to zero). We opted for using the maximum COPE rather than the maximum $t$-statistic because the variance of the response at different time points was very different. Looking up the percentile at which the observed COPE (without sign-flip permutation) occurs gives us our $p$-value. We get a $p$-value for each channel and time point. Sufficiently small $p$-values ($< 0.05$) are deemed to be significant.

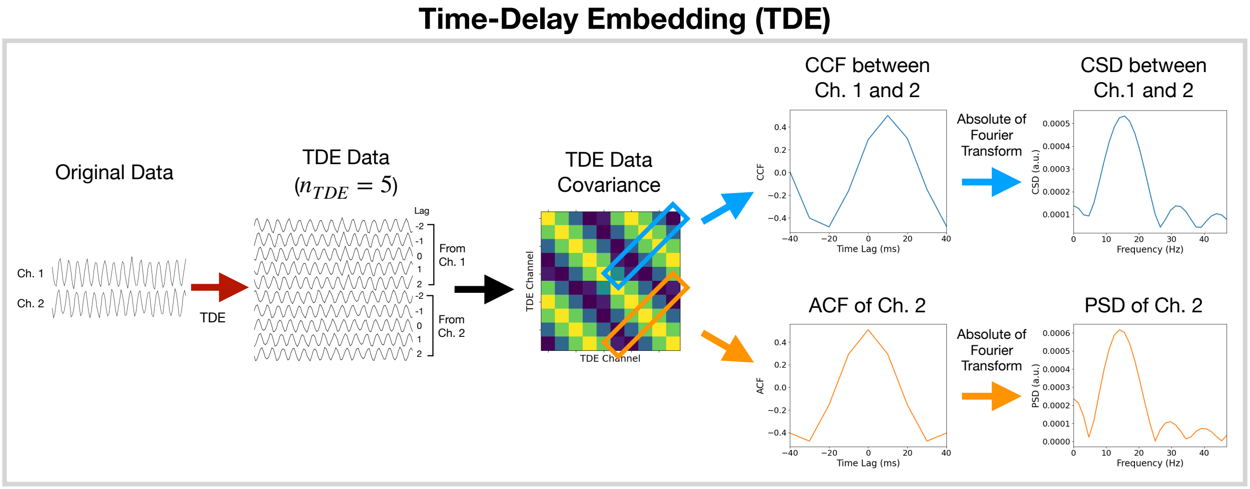


Figure S1: **TDE encodes spectral information (P/CSD via the A/CCF) into the covariance matrix of the TDE data.** Acronyms: time-delay embedding (TDE); power/cross spectral density (P/CSD); auto/cross correlation function (A/CCF); channel (Ch).

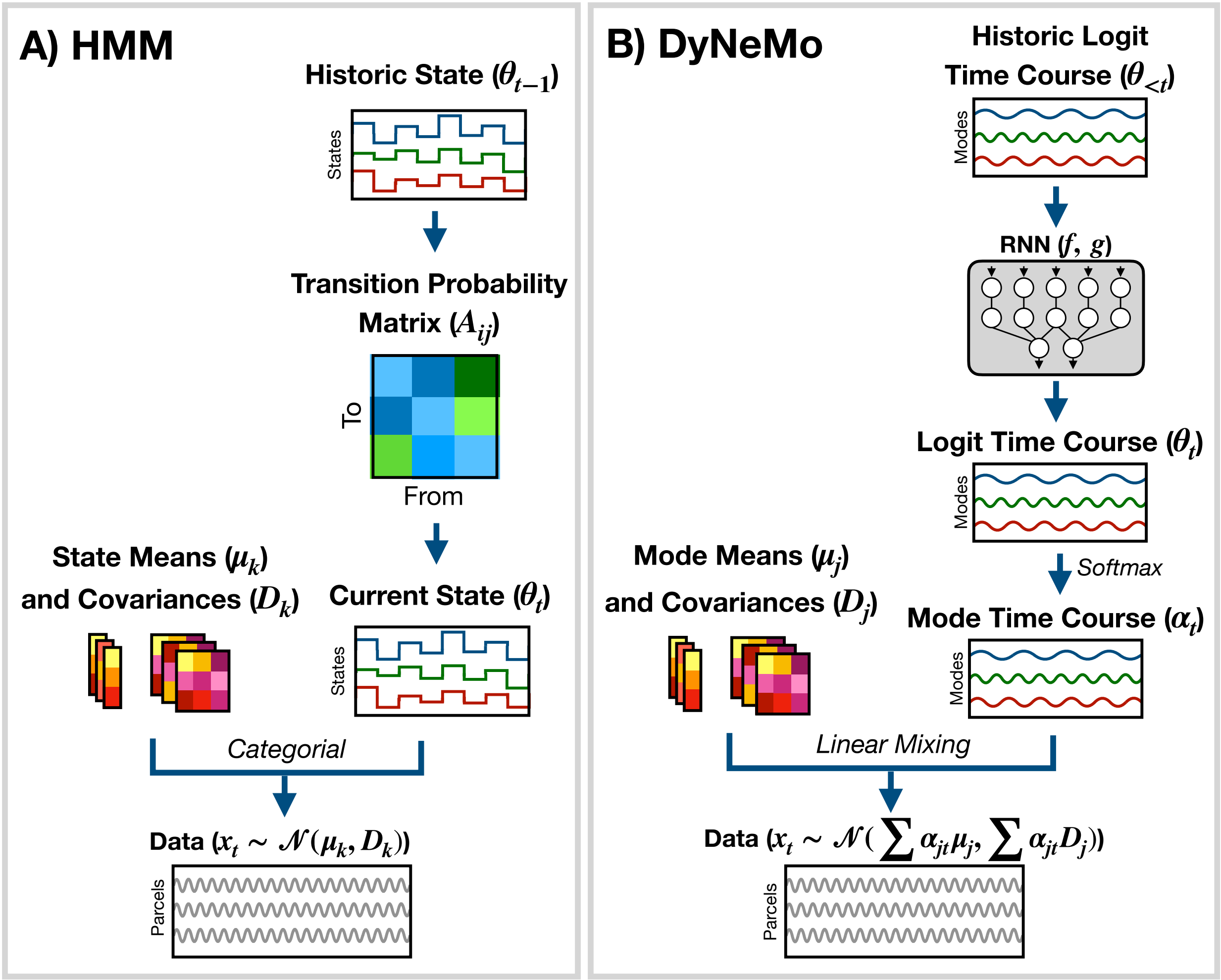


Figure S2: **Overview of dynamic network generative models.** A) Hidden Markov Model (HMM). Here, the data is generated using a hidden state ($\boldsymbol{\theta}_t$) and observation model, which in our case is a multivariate normal distribution parameterised by a state mean ($\boldsymbol{\mu}_k$) and covariance ($\boldsymbol{D}_k$). Only one state can be active at a given time point. Dynamics are modelled via state switching using a transition probability matrix ($\boldsymbol{A}_{ij}$), which forecasts the probability of the current state based on the previous state. B) Dynamic Network Modes (DyNeMo). Here, the data is generated using a linear combination of modes ($\boldsymbol{\mu}_j$ and $\boldsymbol{D}_j$) and dynamics are modelled using a recurrent neural network (RNN: $f$ and $g$), which forecasts the probability of a particular mixing ratio ($\boldsymbol{\alpha}_t$) based on a long history of previous values via the underlying logits ($\boldsymbol{\theta}_t$). Crucially, DyNeMo models the data as a overlapping linear mixture of networks.

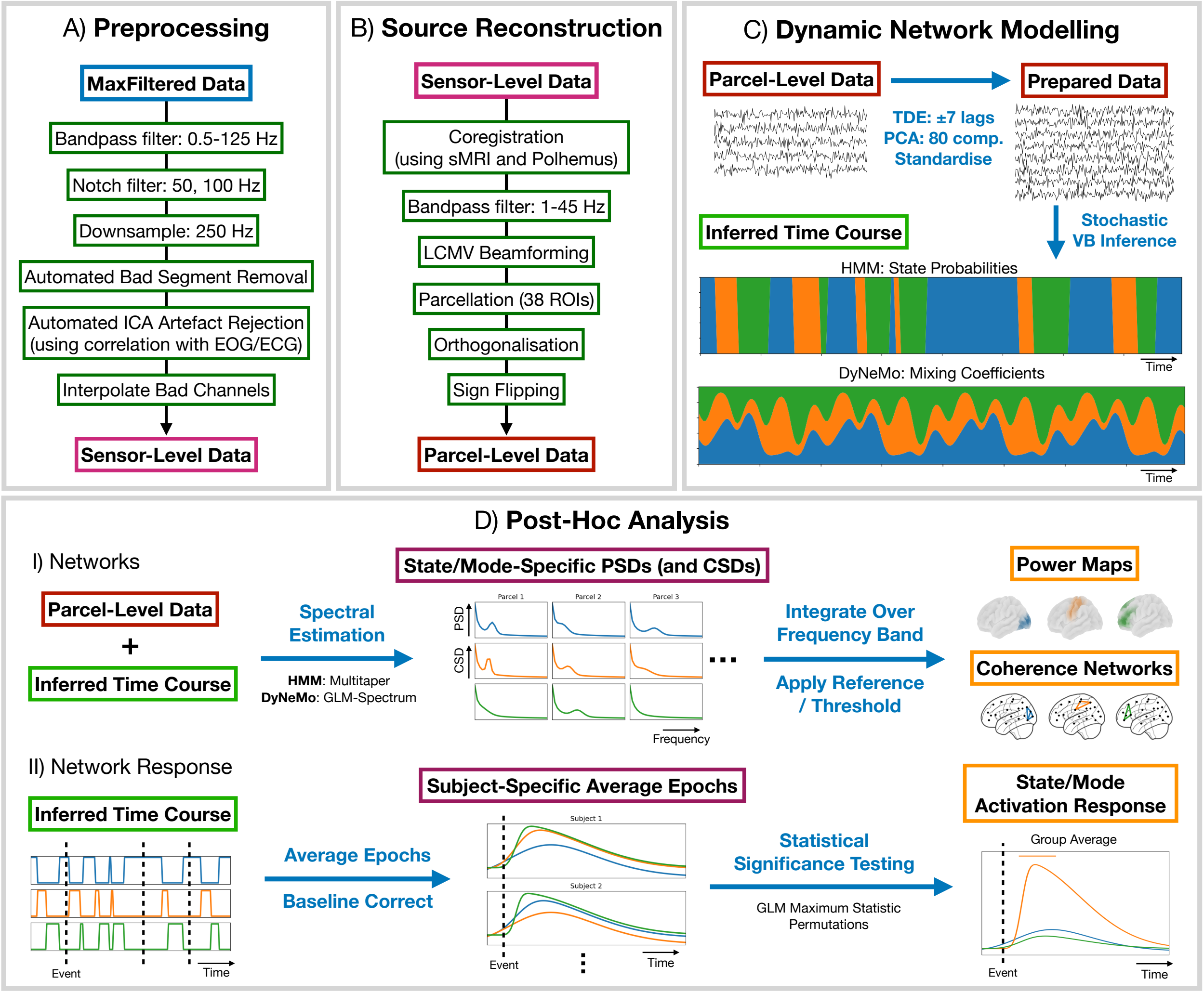


Figure S3: **Start-to-end overview of the dynamic network analysis pipeline.** A) Preprocessing applied to the publicly available raw (MaxFiltered) data. B) Source reconstruction using an LCMV beamformer. C) Dynamic network modelling: this includes first preparing the data then inferring the parameters of our generative model (either the HMM or DyNeMo). D) Post-hoc analysis: I) we use the parcel-level data and inferred state/mode times courses to estimate state/mode specific spectra, which are then used to calculate power maps and coherence networks; II) we use the inferred state/mode time courses to assess which networks show a statistically significant response. Acronyms: Independent Component Analysis (ICA); Electrooculography (EOG); Electrocardiograph (ECG); structural Magnetic Resonance Image (sMRI); Linearly Constrained Minimum Variance (LCMV); Region of Interest (ROI); Time-Delay Embedding (TDE); Principal Component Analysis (PCA); Variational Bayes (VB); Hidden Markov Model (HMM); Dynamic Network Modes (DyNeMo).

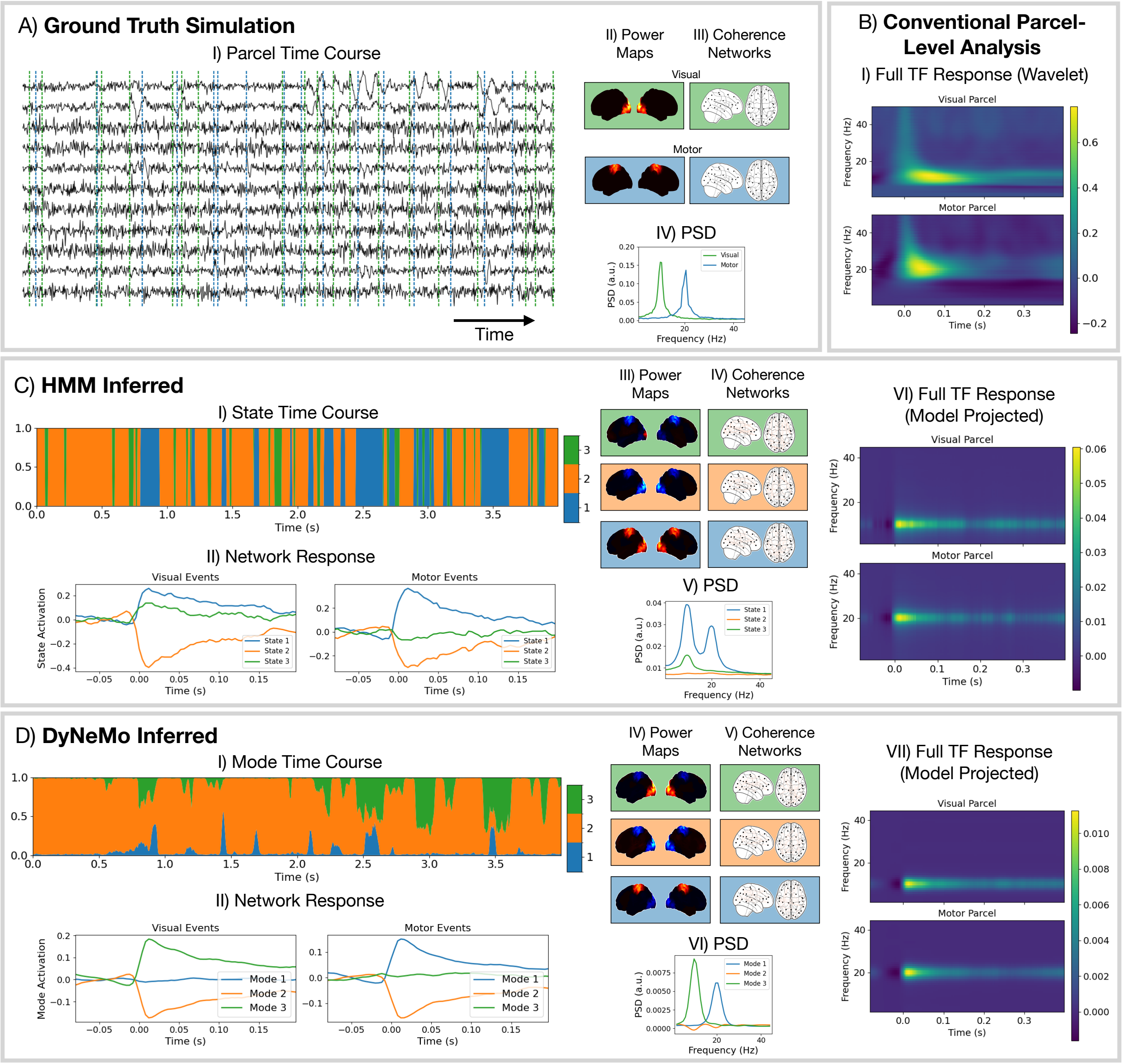


Figure S4: **Simulation without phase coupling.** Reproduction of Figure 4 replacing the simulation of a sine wave with narrowband filtered white noise: 9-11 Hz for the visual network and 19-21 Hz for the motor network. We see the conventional analysis and dynamic network inference are unaffected by the lack of phase coupling and neither dynamics network model erroneously identifies any connectivity.

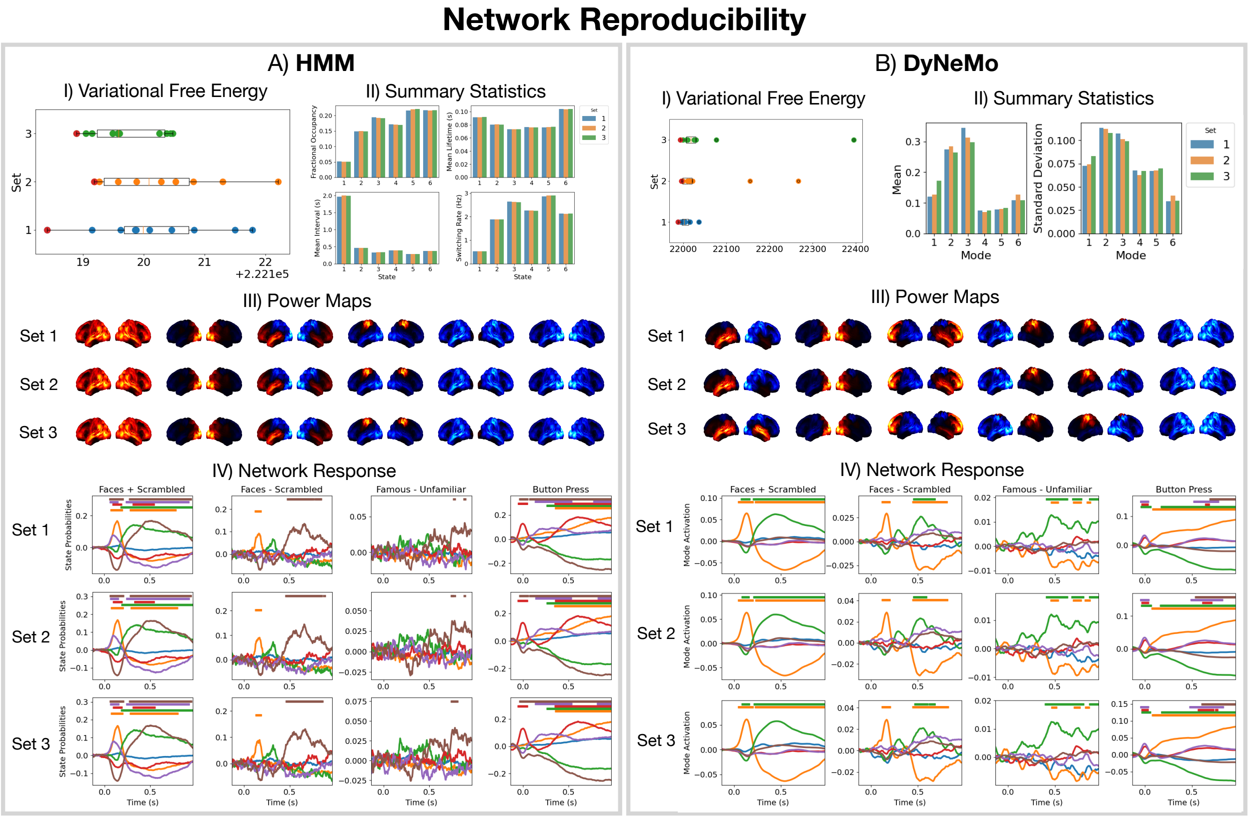


Figure S5: **Reproducibility of inferred networks.** For the Hidden Markov Model (HMM, A) and Dynamic Network Modes (DyNeMo, B): I) variational free energy for 3 sets of 10 runs; II) summary statistics averaged over subjects for the best run (lowest variational free energy) from each set; III) lateral views of power maps relative to the mean across states/modes for the best run from each set; IV) baseline corrected network response for the best run from each set. The horizontal bar indicates a $p$-value $< 0.05$.

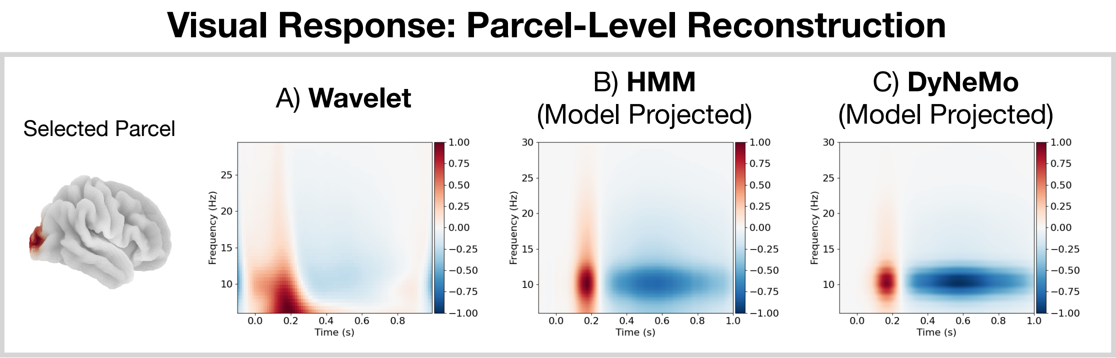


Figure S6: **Full time-frequency (TF) response to visual stimuli.** For the selected parcel, shown on the left: A) wavelet transform of the average parcel time course epoched around visual stimuli; B) TF response of the parcel according to the Hidden Markov Model (HMM) fit calculated by multiplying the baseline corrected state probabilities with the state-specific multitaper spectra and summing. C) TF response of the parcel according to the Dynamic Network Modes (DyNeMo) fit calculated by multiplying the baseline correct mode activations with the mode-specific regression spectra and summing.

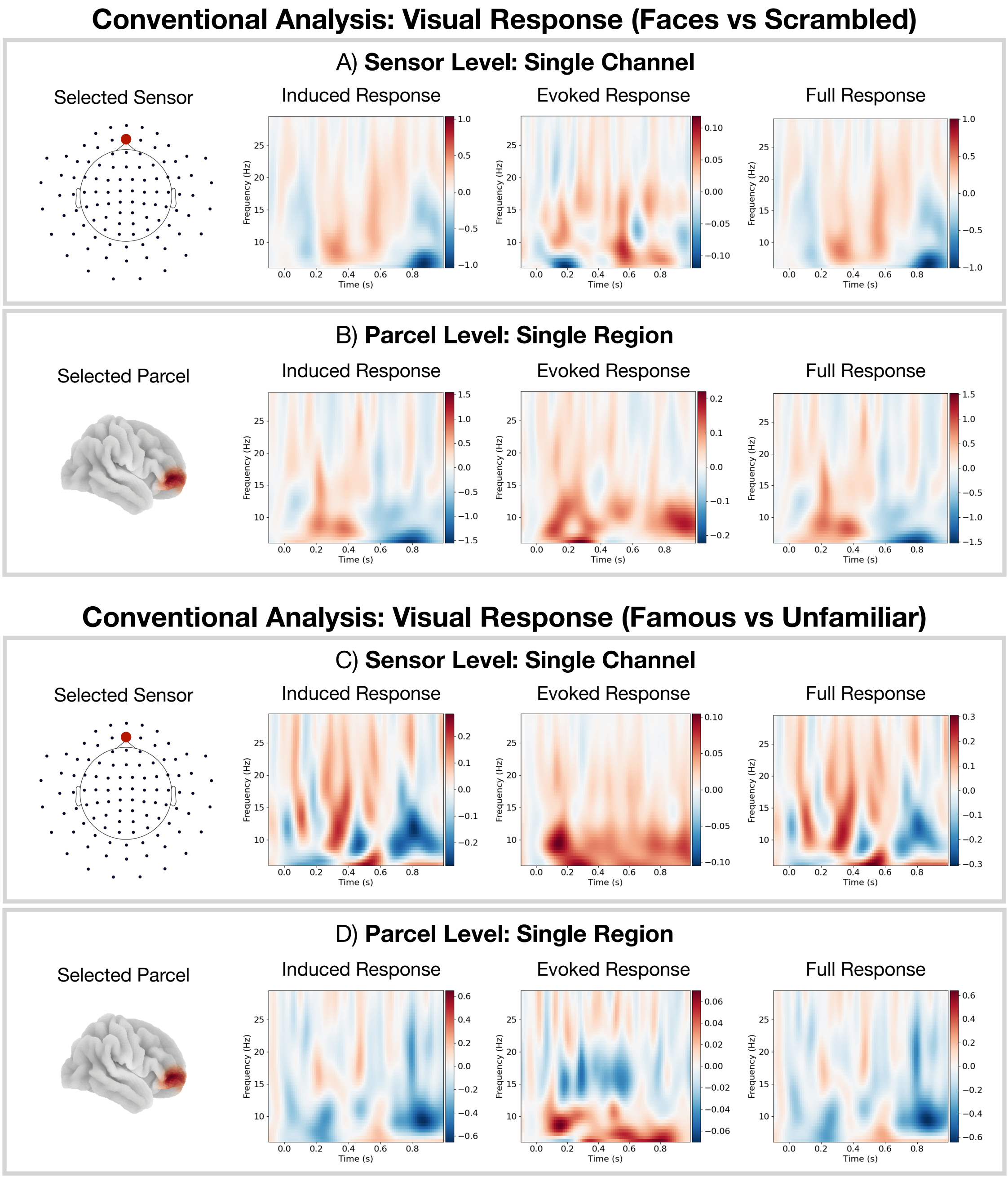


Figure S7: **Conventional time-frequency analysis for a frontal sensor/parcel.** A and B) Faces vs scrambled contrast. C and D) Famous vs unfamiliar faces contrast. There are no significant frequencies or time points.

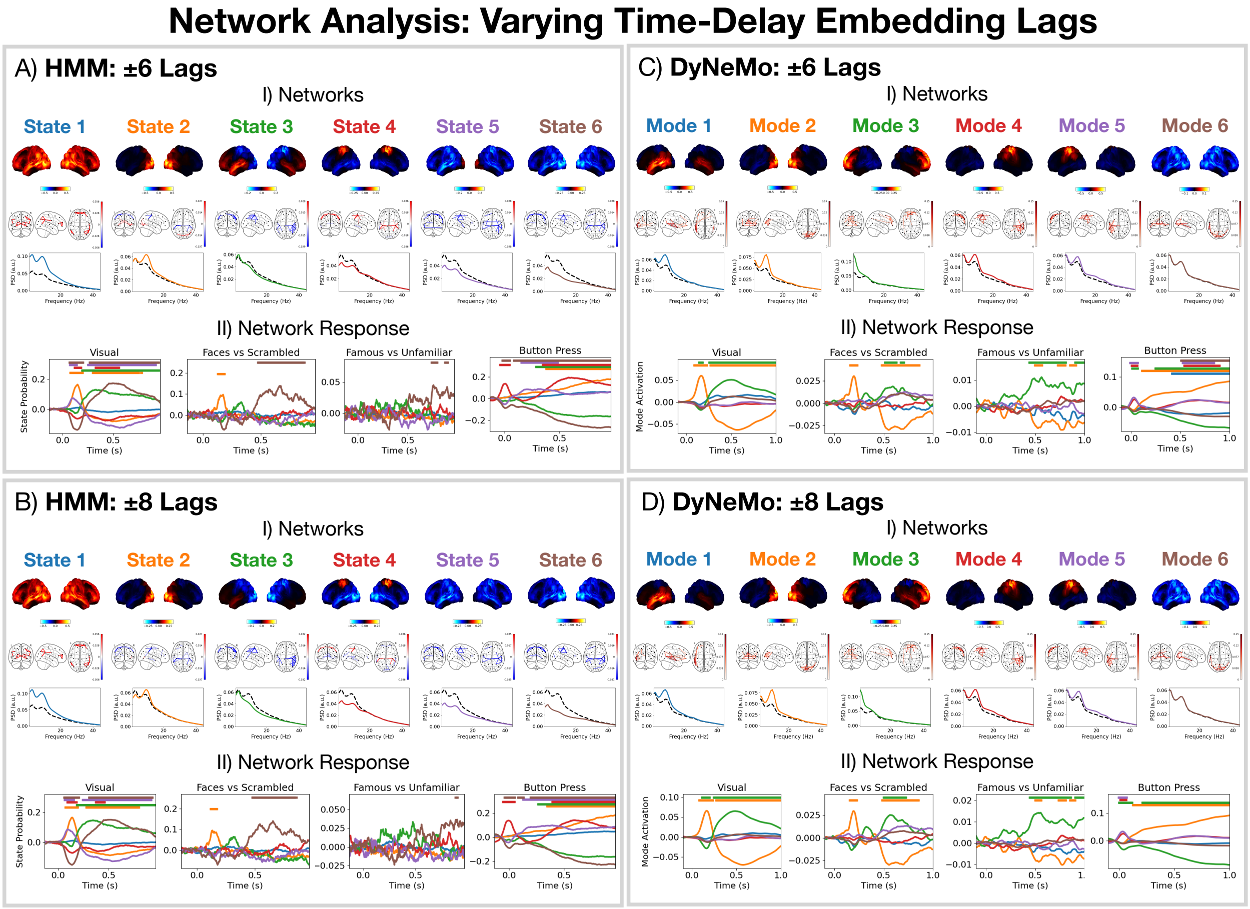


Figure S8: **Effect of a different number of lags in the time-delay embedding.** For the HMM (A and B) and DyNeMo (C and D) power maps, coherence networks and PSDs (I) and network responses for different events (II). Lateral views of the power maps relative to the mean across states/modes from the best of 10 runs are shown. Only the top 2% of edges are shown in the coherence networks. The solid line shows the state/mode specific PSD and the dashed line shows the static PSD. The horizontal bars in each network response indicate time points with $p$-value $< 0.05$.
